# Synaptotagmin-7 is required for synchronous but not asynchronous facilitation of glutamate release at cortical boutons

**DOI:** 10.64898/2025.12.19.695358

**Authors:** Dimitris Kotzadimitriou, Helen Langley, Eleanor McGowan, Philipe R. F. Mendonca, Erica Tagliatti, Yulia Timofeeva, Shyam S. Krishnakumar, Kirill E. Volynski

**Author notes:** These authors contributed equally (Dimitris Kotzadimitriou, Helen Langley).

## Abstract

Short-term synaptic plasticity and the kinetics of neurotransmitter release vary widely across synapses, but population measurements obscure the mechanisms that generate this diversity. While the Ca^2+^ sensor Synaptotagmin-7 (Syt7) has been implicated in facilitation, vesicle replenishment and asynchronous vesicle exocytosis, its precise contributions to these processes remain debated. We used quantal-resolution imaging to measure synchronous and asynchronous glutamate release at individual cortical boutons in wild type and Syt7^-/-^ neurons. Stratifying boutons by release efficacy and applying failure-based analysis to isolate trials where the first action potential evoked no release allowed us to separate facilitation from vesicle depletion. Syt7 deletion selectively eliminated activity-dependent facilitation of synchronous release but left facilitation of asynchronous release intact, although its overall magnitude was reduced. We further show that synchronous and asynchronous events arise from functionally distinct vesicle populations. These findings demonstrate that activity-dependent facilitation of synchronous and asynchronous exocytosis are mechanistically separable, enabling synapses to independently tune distinct temporal components of neurotransmission.

## Introduction

Short term synaptic plasticity enables neurons to encode temporal information and detect activity patterns, yet the molecular mechanisms underlying these computations remain incompletely understood (Abbott and Regehr 2004). It is well established that presynaptic terminals operate within two distinct Ca^2+^ signalling regimes (Eggermann et al. 2011). Fast, high amplitude local Ca^2+^ transients in the tens of micromolar range arise when presynaptic voltage-gated Ca^2+^ channels open during an action potential, generating nano- or microdomains in the active zone that drive synchronous neurotransmitter release within milliseconds through the low affinity sensors Synaptotagmin-1 (Syt1), Synaptotagmin-2 (Syt2) and Synaptotagmin-9 (Syt9) (Xu, Mashimo, and Sudhof 2007). In contrast, slower global residual Ca^2+^ signals in the low micromolar range accumulate during repetitive activity and persist for tens to hundreds of milliseconds to support use-dependent facilitation and delayed asynchronous release. The high affinity Ca^2+^ sensor Synaptotagmin-7 (Syt7) has emerged as a key regulator of these slower presynaptic processes. It binds Ca^2+^ with high affinity and exhibits slow dissociation kinetics enabling it to respond to residual Ca^2+^ long after the synchronous Ca^2+^ transient has decayed (Sugita et al. 2001; Hui et al. 2005; Jackman and Regehr 2017).

Syt7 has been implicated in three major processes: facilitation of vesicular release probability, mediation of asynchronous vesicle fusion, and Ca^2+^-dependent vesicle replenishment (Jackman et al. 2016; Turecek and Regehr 2018; Liu et al. 2014; Huson and Regehr 2020; Bacaj et al. 2013). However, determining which role dominates, or whether they coexist, remains challenging. Some studies show complete loss of facilitation in the absence of Syt7, others report primarily reduced asynchronous release, and several suggest deficits in vesicle pool replenishment (Chen et al. 2017; Wu et al. 2024; Vevea et al. 2021; Jackman et al. 2016) This variability likely reflects synapse-specific differences in active zone structure, Ca^2+^ channel / release sites coupling, and simultaneous expression of multiple Ca^2+^ sensors including Doc2α/β and Syt3 (Weingarten et al. 2022; Weingarten et al. 2024; Wu et al. 2024; Huson and Regehr 2020).

Resolving the function of Syt7 has been hindered by methodological limitations. Population recordings average across heterogeneous synapses, masking facilitation at low efficacy boutons by depression at high efficacy boutons that dominate the combined response. Paired-pulse ratios, the standard readout of short-term plasticity, conflate changes in release probability with changes in vesicle availability caused by depletion. Population approaches also cannot separate synchronous and asynchronous components at individual boutons, forcing reliance on indirect measures such as the asynchronous release fraction. Because this fraction is normalised to total release, changes in synchronous output can alter it even when asynchronous release is unchanged, and synchronous and asynchronous components can vary together in ways that keep the fraction constant despite real changes in either one.

To overcome these limitations, we employed single-bouton quantal imaging using the glutamate sensor SF-iGluSnFR.A184V to monitor synchronous and asynchronous glutamate exocytosis across hundreds of cortical synapses (Marvin et al. 2018; Mendonca et al. 2022). This approach provides unique advantages. Because boutons along a single axon originate from the same presynaptic neuron, they are expected to share broadly similar molecular components of the vesicular release machinery and experience comparable presynaptic action potential waveforms. This design therefore minimises sources of variability that complicate the interpretation of Syt7 effects when comparing synapses of different neuronal types. At the same time, boutons supplied by a single axon exhibit wide heterogeneity in release efficacy, asynchronous output, and short-term plasticity. This allows us to examine how the impact of Syt7 on activity-dependent plasticity and asynchronous release depends on overall vesicular release probability - a question that is difficult to resolve using population-averaged electrophysiology. Our key methodological innovation is a failure-based analysis that examines paired-pulse responses conditional on whether the first stimulus elicited glutamate release. By comparing trials where the first pulse succeeded in triggering exocytosis (promoting depletion of docked vesicles) versus those that failed (no depletion), we could separate true facilitation from depletion-dependent effects on the second response.

We found that Syt7 does not affect basal synchronous release but is essential for facilitation of the synchronous component. In contrast, Syt7 contributes approximately half of asynchronous output yet is dispensable for its facilitation. Efficacy-based stratification combined with failure-based analysis revealed that synchronous and asynchronous release represent functionally segregated pathways governed by distinct rules of short-term plasticity. These findings help resolve apparent discrepancies in the Syt7 literature and provide a framework for understanding how synapses independently regulate distinct temporal components of neurotransmission.

## Results

### Quantal imaging of glutamate release and short-term plasticity at single cortical boutons

To quantify and compare the kinetics and short-term plasticity of vesicular release in wild type and Syt7^-/-^ neurons, we used a previously established imaging approach that resolves quantal glutamate release with millisecond precision across presynaptic boutons supplied by a single axon (Mendonca et al. 2022; Mendonca et al. 2024). The overall experimental design is shown in Fig. 1a. SF-iGluSnFR.A184V (hereafter SF-iGluSnFR) was sparsely expressed in neocortical neurons cultured from wild type or Syt7^-/-^ newborn mice. Whole-cell patch clamp recordings from transfected pyramidal neurons were used to evoke action potentials, while axonal SF-iGluSnFR fluorescence signals reported glutamate release at several tens of boutons per neuron (4 ms resolution). Each recording included nine paired-pulse stimuli at 20 Hz (with 10 s intervals between sweeps for recovery) followed by a 50-pulse train at the same frequency (Fig. 1b). Including the first two action potentials of the train yielded ten paired-pulse measurements per bouton, allowing assessment of short-term plasticity during both paired-pulse and sustained activity.

**Figure 1.**
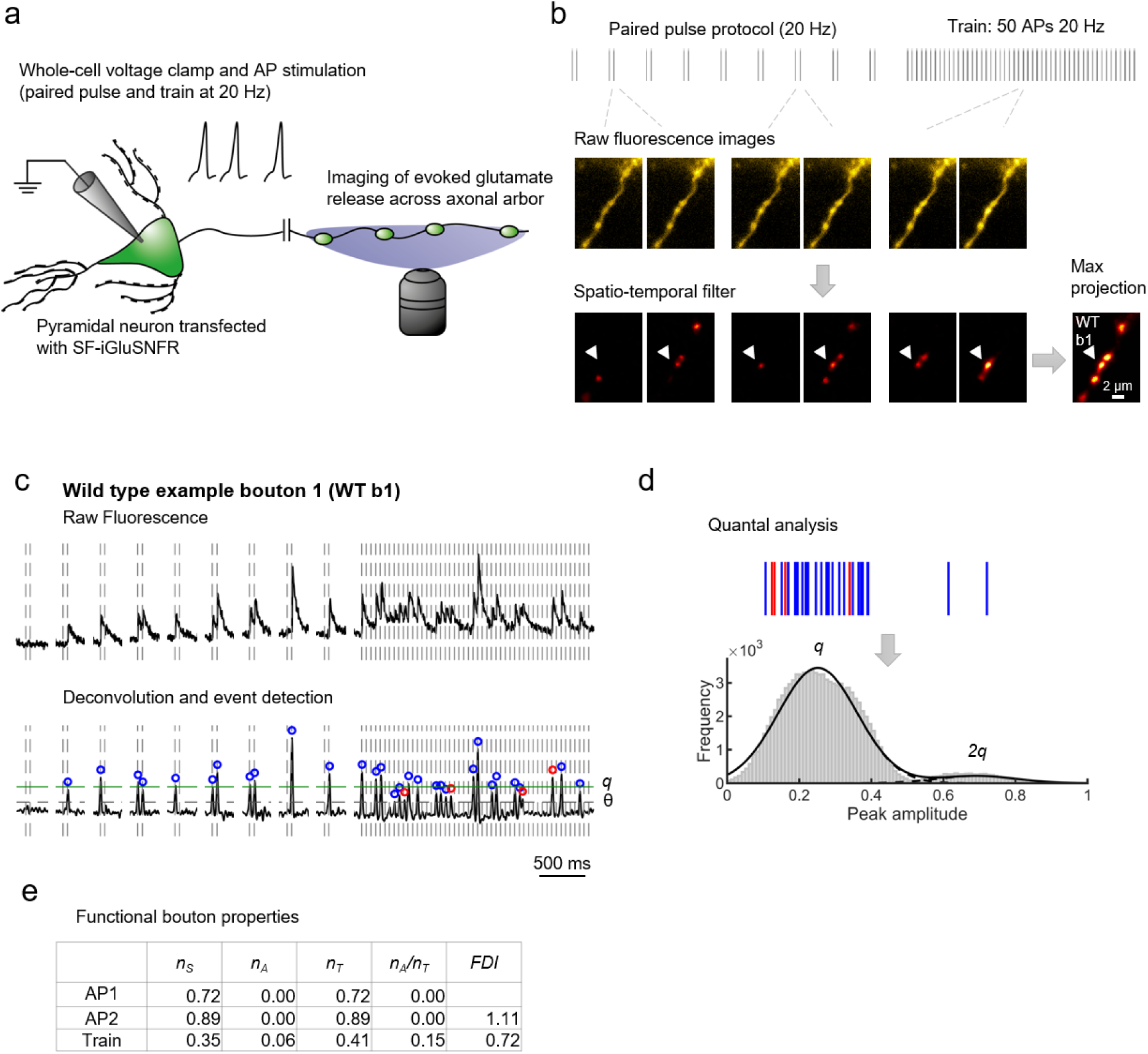
SF-iGluSnFR imaging of quantal glutamate release in individual presynaptic boutons of the axonal arbour. (a) Experimental design. Schematic of the imaging configuration. Whole-cell voltage-clamp recordings were used to evoke action potentials (APs) in SF-iGluSnFR-transfected pyramidal neurons while monitoring glutamate release from their axonal boutons. (b) Identification of active boutons. Image analysis from an axonal fragment of a representative wild type (WT) neuron. Top: stimulation paradigm showing the consecutive paired-pulse (20 Hz) and train (50 APs at 20 Hz) protocols. Example raw fluorescence images acquired after AP stimulation during paired-pulse trials and the train. Bottom: corresponding images after application of a spatiotemporal filter, illustrating vesicular release successes and failures in active boutons within the selected region of interest. Bottom right: maximal projection of the filtered stack, visualising all boutons that released at least one vesicle during the stimulation protocol (five active boutons in this example). (c) Automatic event detection. Top: raw fluorescence trace from an example bouton (b1, WT) indicated by an arrowhead in (b). Bottom: corresponding deconvolved trace showing detected quantal release events. Vertical dotted lines mark AP timings; gaps between traces correspond to 10 s intersweep intervals. Events were identified as local maxima exceeding the detection threshold θ (horizontal dashed line), defined as 4 standard deviations (σ) of baseline fluorescence noise (see Methods). Blue and red circles denote synchronous and asynchronous events (<10 ms and >10 ms after the preceding AP, respectively). (d) Quantal analysis. Amplitudes of all detected events (top raster) were used to generate a quasi-continuous amplitude distribution by bootstrap resampling with bouton specific noise, which improved convergence and stability of the Gaussian mixture fit (see Methods). The resulting distribution was fitted with a sum of Gaussian functions to estimate the quantal amplitude (q, green line on the deconvolved traces in (c)), corresponding to the mean single vesicle SF iGluSnFR response. Peaks at q and 2q indicate one and two vesicle release events, respectively. See additional examples in Supplementary Figs. 1 and 2. (e) Functional bouton parameters for the example bouton b1 (WT). *n_T_*, *n_S_*, and *n_A_* represent total, synchronous, and asynchronous release efficacies (average number of quanta released per action potential); *n_A_*/*n_T_* denotes the asynchronous release fraction, and *FDI*is Facilitation-Depression Index. Parameters were calculated for the 1^st^ (AP1) and 2^nd^ (AP2) stimuli of the paired-pulse protocol and for the steady-state phase of the train (Train; APs 11– 50). See Methods for exact definitions.

Image stacks were analysed using our established analysis pipeline (Fig. 1b–c) (Methods) (Mendonca et al. 2022). After applying spatiotemporal filtering to remove baseline drift, maximal projection images were generated. This allowed us to visualise all active boutons across the axonal arbour that released at least one vesicle during the stimulation protocol. Regions of interest corresponding to active boutons were automatically selected, and fluorescence traces were extracted. Release events were then detected by time-deconvolution of the SF-iGluSnFR signal, which provided the amplitude and timing of individual vesicular release events.

We classified synchronous and asynchronous events using a 10 ms threshold relative to the somatic action potential that was adapted from our previous study (Mendonca et al. 2022). This threshold accounts for axonal conduction delays of approximately 2–6 ms between the soma and boutons (Sabater, Rigby, and Burrone 2021), which blur the temporal distinction between the two release modes. In (Mendonca et al. 2022)., latency distributions were compared between wild type neurons and Syt1^-/-^ neurons that lack synchronous release. Wild type boutons showed a sharp early peak in event latency within the 10 ms window, whereas this peak was largely absent in Syt1^-/-^ neurons, confirming that a 10 ms boundary provides a conservative separation between synchronous and asynchronous release. This approach minimises false classification of synchronous events as asynchronous, although it may underestimate the asynchronous component.

Amplitude distributions of deconvolved fluorescence transients at individual boutons typically showed discrete peaks consistent with the release of one or more vesicles (Fig. 1d and Supplementary Figs. 1 and 2). Quantal amplitude (q) was estimated by fitting these distributions with a sum of Gaussian functions (see Methods). From these data we derived bouton level measures of vesicular release for each part of the stimulation protocol. For the paired-pulse protocol, we defined the total release efficacy at the first and second action potentials as *n_T_* (1) and *n_T_* (2), calculated as the mean number of quanta released per action potential across the ten paired-pulse trials. For the train protocol, the steady-state total release efficacy *n_T_* (*Tr*) was defined as the mean number of quanta released per action potential across action potentials 11 to 50. In the same way, we obtained synchronous and asynchronous release efficacies *n_S_* (1), *n_S_* (2), *n_S_* (*Tr*) and *n_A_* (1), *n_A_* (2), *n_A_* (*Tr*). From these values we then calculated the asynchronous release fraction *n_A_* /*n_T_* for each part of the protocol (Fig. 1e and Supplementary Figs. 1 and 2).

**Figure 2.**
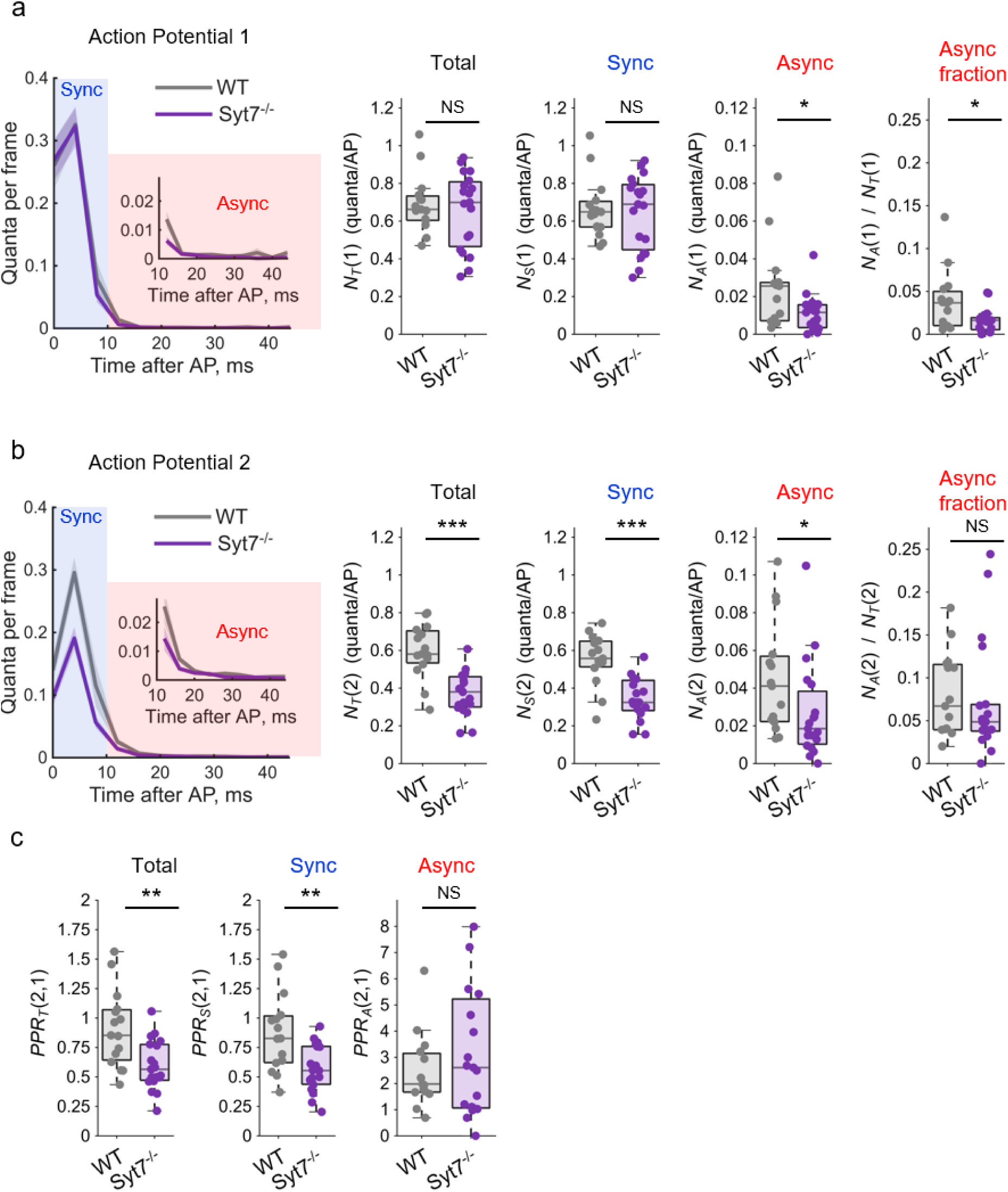
Cell level analysis of glutamate release in Syt7^-/-^ and wild type neurons during paired-pulse protocol. (a, b) Quantification of vesicular release at the first (a) and second (b) action potentials. Left panels: average time courses of evoked glutamate release in wild type (WT) and Syt7^-/-^ neurons (quanta per frame). For each bouton, the number of detected quanta in individual frames following the first or second action potential was averaged across ten paired-pulse trials (frames 0–11; 4 ms per frame). Bouton responses were then averaged within each neuron and subsequently across neurons (shaded areas indicate SEM across cells). A 10 ms threshold after the somatic action potential (between frames 2 and 3, i.e. at 10 ms) was used to separate synchronous and asynchronous events (see Methods). Right panels: cell-averaged total, synchronous, asynchronous release, and the asynchronous fraction. (c) Paired-pulse ratios for total, synchronous, and asynchronous components, calculated as the ratio of cell-averaged release at the second versus the first action potential. One Syt7^-/-^ neuron lacked asynchronous release on the first pulse in all boutons but exhibited asynchronous events on the second; because the corresponding PPR is undefined, this datapoint is not plotted but was treated as maximally facilitated when computing the median across cells. Boxplots show the median and interquartile range with whiskers extending to 1.5x the interquartile range and individual datapoints overlaid. Wild type, n = 15 cells; Syt7^-/-^, n = 19 cells. * p < 0.05, ** p < 0.01, *** p < 0.001, NS not significant; two-tailed Mann–Whitney U tests. Exact p values and additional statistical details are provided in Supplementary Table 1 and SourceData.xlsx.

We also quantified short-term synaptic plasticity in each bouton. The conventional paired-pulse ratio (*PPR*), defined as the ratio of the test response to the response at the first pulse, is suitable for cell level analysis because averaging across many boutons ensures that the first stimulus almost always produces a measurable response. At the level of individual boutons, however, zero release at the first action potential is common in low efficacy synapses, which makes the conventional *PPR* unstable or undefined. To obtain a measure that remains well defined in all boutons, we used the Facilitation-Depression Index (*FDI*), which normalises the test response to the mean of the first and test responses and therefore avoids division by zero. For the paired-pulse protocol, *FDI_T_* (2,1) was calculated as *n_T_* (2) divided by *mean* [*n_T_* (1), *n_T_* (2)], thereby comparing the second response with the first. Similarly, for the train protocol we calculated *FDI_T_* (*Tr*,1) as *n_T_* (*Tr*) divided by *mean* [*n_T_* (*Tr*), *n_T_* (2)], comparing the steady-state with the first response. Similar to *PPR*, *FDI* values above 1 indicate facilitation and values below 1 indicate depression, but because the test response is normalised to the mean of the two responses, the index approaches an upper limit of 2 (see Supplementary Fig. 3).

Presynaptic boutons in both genotypes showed pronounced variability in release efficacy, kinetics, and short-term plasticity (Fig. 1; Supplementary Figs. 1 and 2). Such heterogeneity was evident both across neurons and among boutons supplied by the same axon. During paired-pulse stimulation and at the onset of trains, individual release events corresponded to the fusion of one and sometimes up to five vesicles, whereas during the steady-state phase release became predominantly single-quantal, particularly in boutons with low release efficacy.

Deconvolution resolved the vast majority of release events during 20 Hz trains, although a small proportion of events fell below the detection threshold because of temporal overlap and partial indicator saturation (see Methods). This under-detection may slightly underestimate absolute efficacy values but it affects both genotypes and therefore does not bias relative comparisons.

In summary, this imaging approach provided quantal-resolution measurements of synchronous and asynchronous vesicular release across large bouton populations within single neurons, enabling direct comparison of their kinetics and short-term plasticity between wild type and Syt7^-/-^ synapses.

### Syt7 knockout differentially affects synchronous and asynchronous release at the cell level

To enable comparison with previous electrophysiological and imaging studies, release efficacies were averaged across boutons within each neuron to obtain cell-level values: *N_T_* = 〈*n_T_*〉, *N_S_* = 〈*n_S_* 〉 and *N_A_* = 〈*n_A_* 〉 (capital letters denote cell-averaged responses throughout the manuscript). Recordings were obtained from 15 wild type and 19 Syt7^-/-^ neurons, each contributing between 21 and 177 active boutons (median 59 per neuron). This analysis allowed quantification of genotype-dependent differences in synchronous and asynchronous release during both paired-pulse stimulation (Fig. 2) and sustained 20 Hz trains (Fig. 3). A summary of cell-level results is provided in Supplementary Table 1.

**Figure 3.**
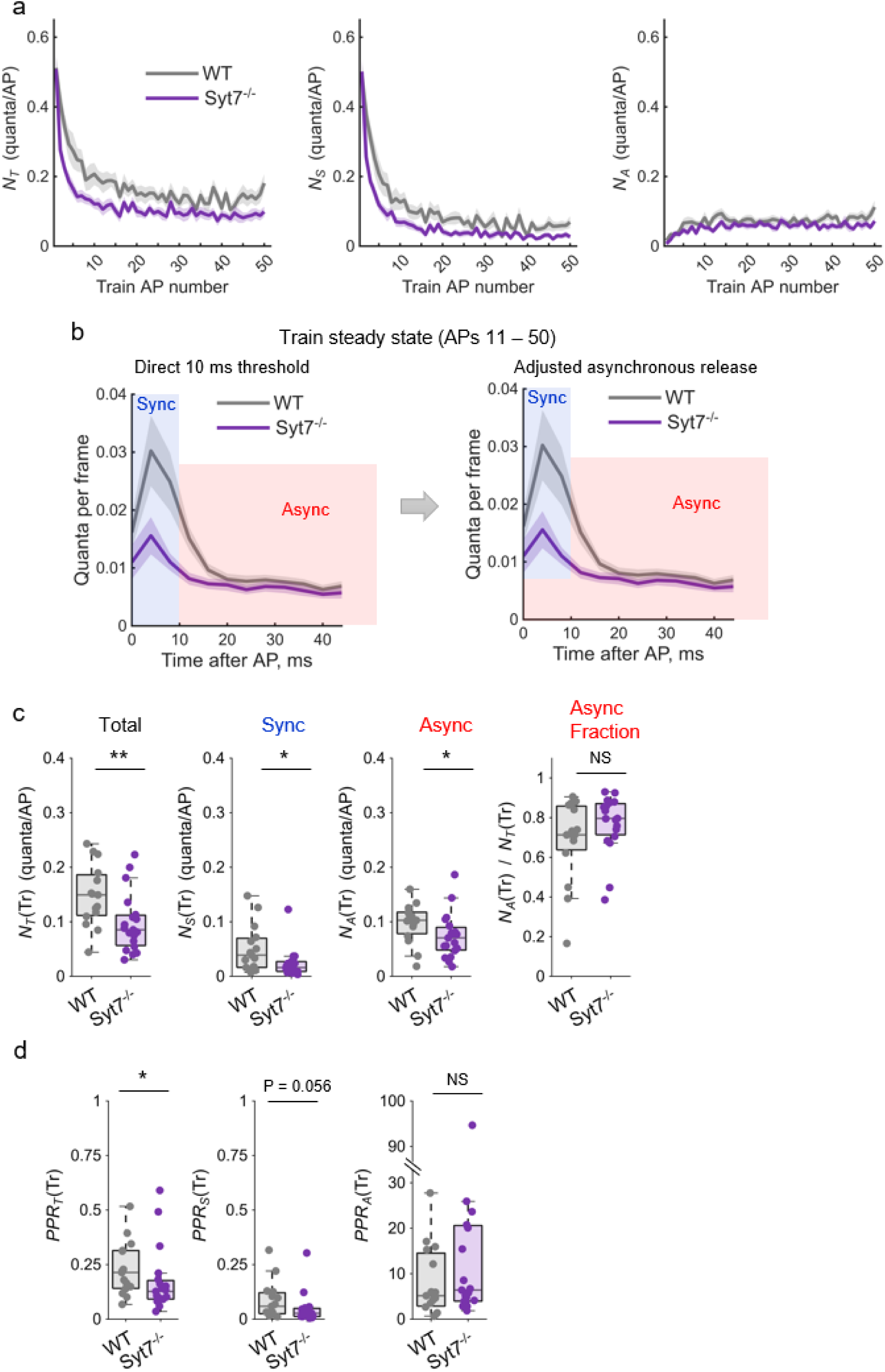
Cell level analysis of glutamate release in Syt7^-/-^ and wild type neurons during the steady-state of the 20 Hz train. (a) Comparison of total, synchronous, and asynchronous vesicular release per action potential in wild type (WT) and Syt7^-/-^ neurons during 20 Hz trains of 50 stimuli. Traces represent cell-averaged vesicular release per bouton for each action potential, shown as the mean across neurons (shaded areas indicate SEM across cells). Synchronous and asynchronous events were separated using a 10 ms threshold after the somatic action potential, as in Fig. 2. (b) Average time courses of evoked glutamate release during the steady-state portion of the train (APs 11–50). Left: classification of synchronous and asynchronous components using a fixed 10 ms threshold. Right: corrected classification in which the portion of ongoing asynchronous release falling within the 10 ms window was reassigned from synchronous to asynchronous bins (see Methods). Shaded areas indicate SEM across cells. (c) Cell-averaged adjusted total, synchronous, and asynchronous release during the steady-state after correction for ongoing asynchronous release, and the corresponding asynchronous fraction. (d) Paired-pulse ratios for total, synchronous and asynchronous release during the train steady-state, calculated as the ratio of the cell-averaged release at APs 11–50 to the release at the first action potential. One Syt7^-/-^ neuron showed no detectable asynchronous release on the first pulse and therefore did not yield a defined asynchronous PPR; this datapoint is not shown and was treated as maximally facilitated when computing the median. Boxplots show the median and interquartile range, with whiskers extending to 1.5× the interquartile range and individual datapoints overlaid. Wild type, n = 15 cells; Syt7-/-, n = 19 cells. * p < 0.05, ** p < 0.01, NS not significant; two-tailed Mann–Whitney U tests. Exact p values and additional statistical details are provided in Supplementary Table 1 and SourceData.xlsx.

During paired-pulse stimulation, most release events were synchronous, with a smaller asynchronous component that decayed rapidly (Fig. 2a, b). At the first action potential (Fig. 2a), total *N_T_* (1) and synchronous *N_S_* (1) release were comparable between genotypes, indicating that under our recording conditions Syt7 knockout does not alter basal release efficacy. In contrast, median asynchronous release efficacy *N _A_* (1) was reduced by approximately 50% in Syt7^-/-^ neurons compared with wild type. Because asynchronous events accounted for less than 4% of total release in both genotypes, this reduction had little effect on overall release efficacy but lowered the asynchronous release fraction *N_A_* (1)/*N_T_* (1).

At the second action potential (Fig. 2b), total *N_T_* (2), synchronous *N_S_* (2) and asynchronous *N _A_* (2) release were all reduced in Syt7^-/-^ neurons compared with wild type (Fig. 2b). For total and synchronous components this reflected marked paired-pulse depression in Syt7^-/-^, with median *PPR_T_* (2,1) = *N_T_* (2)/*N_T_* (1) and *PPR_S_* (2,1) = *N_S_* (2)/*N_S_* (1) values of 0.56, compared to ∼0.83 in wild type (Fig. 2c). Asynchronous release efficacy *N_A_* (2) remained about 50% lower in Syt7^-/-^ neurons than in wild type, but both genotypes showed a comparable relative increase between the first and second stimuli, with median *PPR_A_* (2,1) = *N_A_* (2)/*N_A_* (1) values of 2.0 in wild type and 2.6 in Syt7^-/-^ (Fig. 2c). Because synchronous and asynchronous components decreased in parallel in Syt7^-/-^, the asynchronous release fraction *N_A_* (2)/*N_T_* (2) did not differ significantly between genotypes, illustrating that the fraction is not a reliable indicator of changes in asynchronous output when both components vary simultaneously.

To determine how Syt7 knockout affects vesicular release during sustained repetitive activity, we next analysed responses to 20 Hz trains of 50 action potentials (Fig. 3). In both genotypes, total and synchronous release efficacy declined sharply at the start of the train and reached a steady-state after approximately the tenth stimulus (APs 11–50). This reduction was more pronounced in Syt7^-/-^ neurons. Conversely, the asynchronous component showed strong facilitation in both genotypes, with only a small decrease in its steady-state efficacy of asynchronous release in Syt7^-/-^ neurons compared with wild type (Fig. 3a).

As in the paired-pulse protocol, during the train steady-state we averaged the release time course relative to each action potential across APs 11–50 (Fig. 3b). In both genotypes, the event rate exhibited a synchronous peak within 10 ms followed by a slower tail that remained approximately constant. Because the interstimulus interval at 20 Hz is 50 ms, this late constant asynchronous component continues into the next cycle. A fixed 10 ms boundary therefore misclassifies part of this ongoing asynchronous release as synchronous at the onset of the following action potential, leading to an overestimation of *n_S_* (*Tr*) and underestimation of *n_A_* (*Tr*). To correct this bias and estimate the true synchronous component, we subtracted from the synchronous bin the residual asynchronous contribution of the preceding action potential and reassigned it to the asynchronous bin, as illustrated in Fig. 3b and detailed in the Methods. In all subsequent analysis we used the adjusted release efficacy values but for simplicity retained the same notation as before: *n_S_* (*Tr*) and *n_A_* (*Tr*).

We next compared total, synchronous, and asynchronous release during the steady-state of the train at the cell level (Fig. 3c). Based on median values, total release efficacy *N_T_* (*Tr*) was reduced by approximately 40% in Syt7^-/-^ neurons relative to wild type. The synchronous component *N_S_* (*Tr*) showed a stronger reduction of about 60%, whereas the asynchronous component *N_A_* (*Tr*) decreased by around 30%. Because synchronous release was more strongly affected than asynchronous release, the asynchronous release fraction *N_A_* (*Tr*)/*N_T_* (*Tr*) appeared slightly higher in Syt7^-/-^ neurons (median 0.80 versus 0.71 in wild type), although this difference was not statistically significant. We also calculated the paired-pulse ratios for each release component during the steady-state phase of the train (Fig. 3d). Depression of total and synchronous release was approximately twofold stronger in Syt7^-/-^ neurons compared with wild type. In contrast, asynchronous release showed similar facilitation in both genotypes, increasing by approximately fivefold in wild type and sixfold in Syt7^-/-^.

In summary, Syt7 knockout leaves basal synchronous release unchanged but causes strong depression during repetitive stimulation, consistent with previous reports (Jackman et al. 2016; Liu et al. 2014). Asynchronous release shows a different pattern: although its absolute magnitude is reduced in Syt7^-/-^ neurons under both basal conditions and during trains, facilitation of the remaining asynchronous component is maintained. This dissociation between output magnitude and plasticity is consistent with synchronous and asynchronous facilitation relying on distinct Ca^2+^-dependent pathways.

### Bouton-level analysis of release properties across efficacy classes

Because release kinetics and short-term plasticity vary substantially among boutons supplied by the same axon, we analysed the effect of Syt7 knockout at the level of individual boutons to determine whether genotype-dependent differences were uniform across synapses or influenced by bouton-specific release efficacy. Averaging across boutons within each neuron can obscure such dependencies, as cell-wise averages are dominated by high-efficacy boutons. These boutons are particularly prone to vesicle depletion during repetitive stimulation, which biases cell-averaged responses towards depression. Moreover, asynchronous release is typically lower at high-efficacy boutons (Mendonca et al. 2022), potentially masking Syt7-dependent effects that vary with release efficacy. To address these issues, we examined bouton-level data directly, in order to assess how Syt7 influences release kinetics (Fig. 4) and short-term plasticity (Fig. 5) across boutons differing in release efficacy.

**Figure 4.**
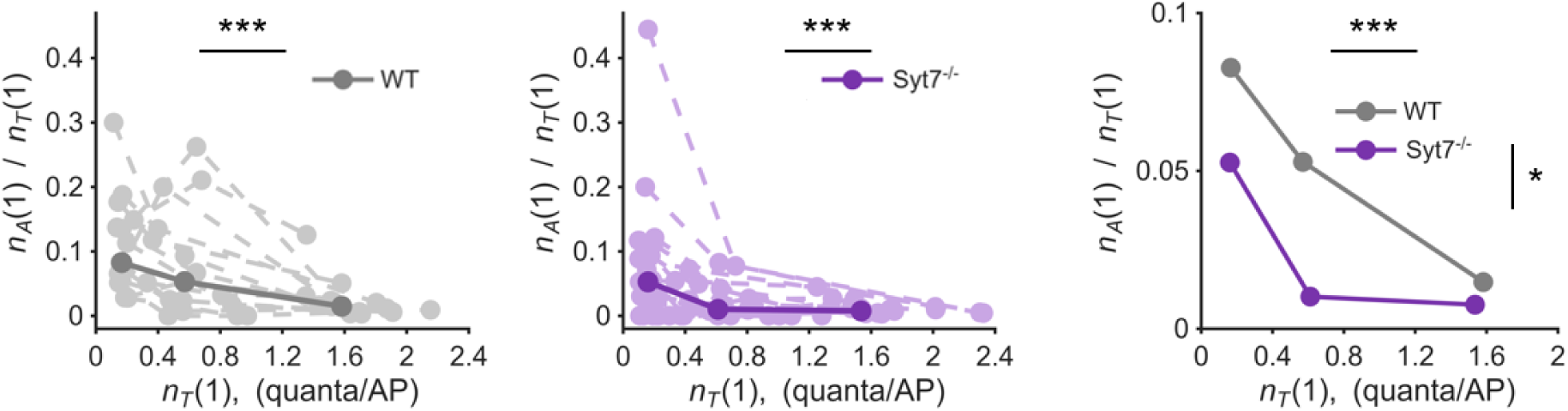
Bouton-level dependence of the asynchronous release fraction on release efficacy. Boutons within each neuron were sorted by total release efficacy at the first action potential *n_T_* (1) and divided into three equal-sized groups (terciles). For each neuron, *n_T_* (1) and the asynchronous release fraction *n_A_* (1)/*n_T_* (1) were averaged within each tercile. Left and middle panels: per-cell, per-tercile values for wild type and Syt7^-/-^, respectively, showing the dependence on release efficacy within each genotype. Dashed lines connect terciles from the same neuron; bold points and solid lines indicate population medians. Right panel: comparison of tercile medians between genotypes, illustrating the main effect of genotype across efficacy classes. Statistical significance was assessed using linear mixed-effects models, with Model 1 testing efficacy dependence within genotype (left and middle panels) and Model 2 testing genotype effects across efficacy classes (right panel); the genotype-by-efficacy interaction was not significant. Wild type, n = 15 cells; Syt7-/-, n = 19 cells. Exact statistics are provided in SourceData.xlsx. Significance bars indicate p values (*** p < 0.001; * p < 0.05).

**Figure 5.**
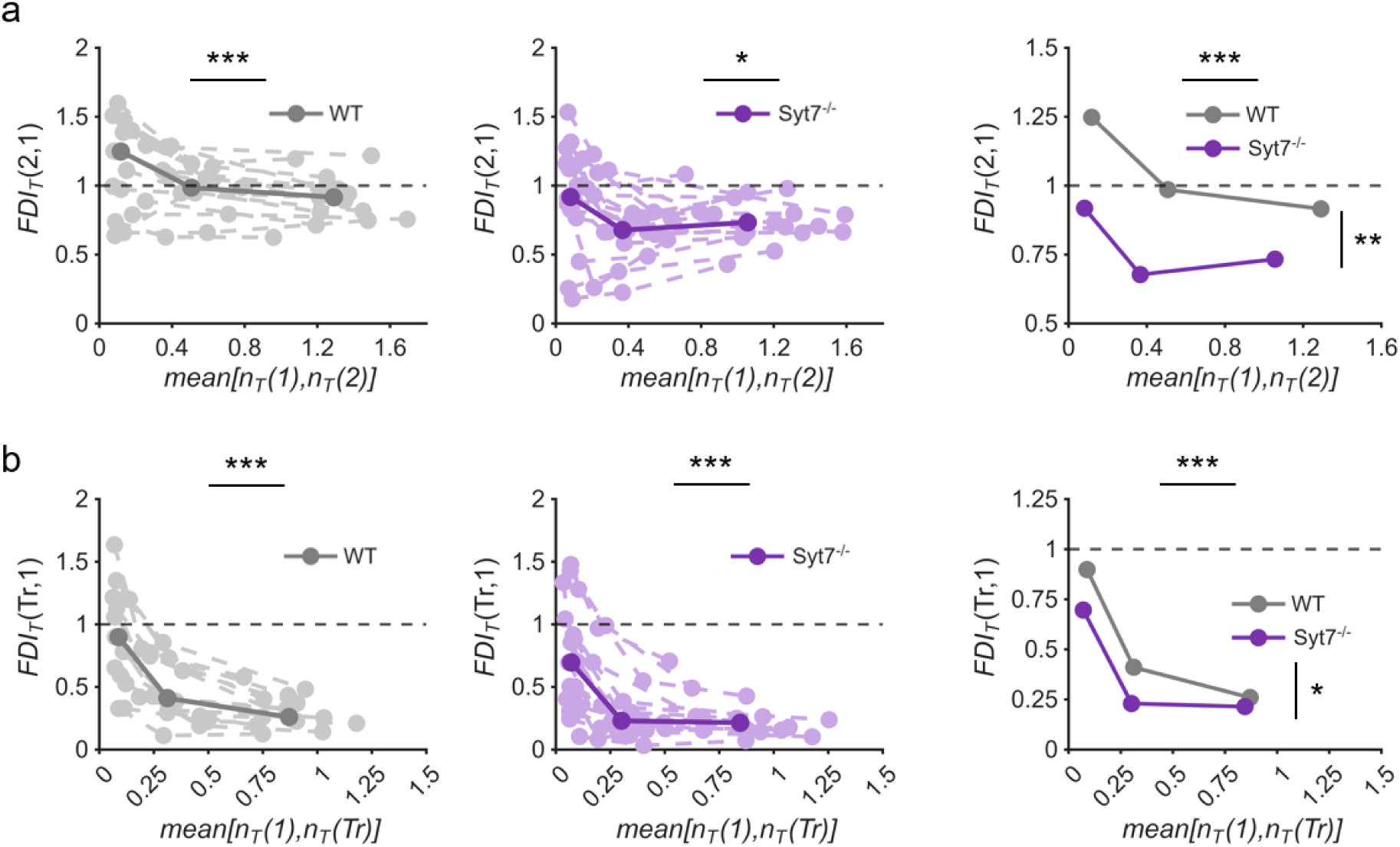
Effect of Syt7 knockout on short-term plasticity across boutons with different release efficacy. Boutons within each neuron were sorted by total release efficacy and divided into three terciles, as in Fig. 4. To avoid bias introduced by sorting on the first-pulse response, grouping was based on the mean total release across the two compared responses. For each neuron and tercile, short-term plasticity was quantified using the Facilitation-Depression Index and plotted against the corresponding tercile mean. (a) Paired-pulse stimulation, *FDI_T_* (2,1). (b) Steady-state phase of the train, *FDI_T_* (*Tr*,1). Left and middle panels: per-cell, per-tercile *FDI* values for wild type and Syt7^-/-^, respectively. Dashed lines connect terciles from the same neuron; bold points and solid lines indicate population medians. Right panels: comparison of tercile medians between genotypes, illustrating loss of facilitation in low-efficacy Syt7^-/-^ boutons and enhanced depression at medium and high efficacy. The dashed horizontal line at *FDI* = 1 indicates no change between responses. Statistical significance was assessed using linear mixed-effects models, with Model 1 testing efficacy dependence within genotype (left and middle panels) and Model 2 testing genotype effects across efficacy classes (right panels); the genotype-by-efficacy interaction was not significant (see Methods). Wild type, n = 15 cells; Syt7-/-, n = 19 cells. Exact statistics are provided in SourceData.xlsx. Significance bars indicate p values (*** p < 0.001; ** p < 0.01; * p < 0.05).

For each selected part of the stimulation protocol, such as the first or second action potential in the paired-pulse paradigm or the steady-state phase of the train, we ranked boutons within each neuron by their total release efficacy *n_T_*, measured during that phase. Following our previous approach (Mendonca et al. 2022), we then divided the boutons into three equal-sized groups (terciles) corresponding to low-, medium-, and high-efficacy classes. For each tercile, we calculated the mean for *n_T_* and for either the asynchronous release fraction or the *FDI*.

This analysis revealed how each parameter varied across boutons of different release efficacy within individual neurons (Supplementary Fig. 4).

To statistically assess these relationships while accounting for cell-to-cell variability, we applied two linear mixed-effects models (LMMs) as described in the Methods. Model 1 tested how each parameter varied with total release efficacy within a genotype, and Model 2 tested for genotype-dependent differences across efficacy classes.

### Syt7 knockout reduces asynchronous release across boutons with high and low release efficacy

To assess the effect of Syt7 knockout on the asynchronous release fraction, we first quantified basal release during the first action potential (Fig. 4). This approach avoided confounds arising from the differential effects of Syt7 knockout on facilitation of synchronous and asynchronous release described above, ensuring that the measured asynchronous fraction *n_A_* (1)/*n_T_* (1) accurately reflected the basal contribution of Syt7 to asynchronous vesicle fusion.

In wild type neurons, *n_A_* (1)/*n_T_* (1) was highest at low-efficacy boutons and declined with increasing efficacy, consistent with our previous work (Mendonca et al. 2022). Syt7^-/-^ neurons exhibited the same dependency but with values uniformly reduced across all terciles. Statistical analysis confirmed a significant main effect of genotype, but no significant genotype-by-efficacy interaction, indicating that Syt7 knockout reduces basal asynchronous release similarly across boutons with different release efficacy.

### Differential effect of Syt7 knockout on short-term plasticity across boutons with different release efficacy

To assess the effect of Syt7 knockout on short-term plasticity at the bouton level, we focused on total release efficacy, since quantifying plasticity separately for synchronous and asynchronous components was not feasible. Many boutons exhibited no detectable asynchronous release throughout the stimulation protocol, which would lead to undefined ratios when calculating component-specific plasticity indices due to division by zero.

Short-term plasticity at individual boutons was quantified using the Facilitation-Depression Index (*FDI*), comparing either the second action potential with the first during paired-pulse stimulation (*FDI_T_* (2,1)) or the steady-state phase of the train with the first action potential (*FDI_T_* (*Tr*,1)). Boutons were ranked by total release efficacy and divided into terciles, as described above. To avoid bias introduced by sorting on the first-pulse response, grouping was based on the average total release across the two compared responses, matching the normalisation used for the *FDI*.

When boutons were compared across efficacy classes, wild type neurons showed pronounced facilitation during paired-pulse stimulation in low-efficacy boutons (median *FDI_T_* (2,1) ≈ 1.25; Fig. 5a). Facilitation progressively declined with increasing release efficacy and converted to depression in high-efficacy boutons, consistent with vesicle depletion of the readily releasable vesicle pool. In contrast, Syt7^-/-^ boutons showed no facilitation even in the low-efficacy group, where depletion effects are minimal (median *FDI_T_* (2,1) ≈ 1.0), and exhibited stronger depression at medium and high efficacy. This pattern is consistent with Syt7 supporting paired-pulse facilitation independently of vesicle depletion, as its loss abolishes facilitation even in boutons where depletion is minimal.

During the steady-state of the train, release reflects the interplay between vesicle depletion, replenishment, and facilitation of vesicular release probability. Under these conditions, both wild type and Syt7^-/-^ boutons showed depression across all efficacy classes (Fig. 5b). Depression was more pronounced in boutons with higher release efficacy, consistent with greater depletion of readily releasable vesicles. Statistical analysis using Model 2 confirmed that Syt7 knockout enhanced the extent of depression, consistent with the cell-level results. This effect was limited in boutons with low and medium release efficacy and was absent in high-efficacy boutons, where both genotypes showed comparable depression. This pattern is inconsistent with a primary role for Syt7 in vesicle replenishment, which would predict the largest genotype difference where depletion is greatest, though our statistical models were not optimised to formally test this interaction.

### Failure-based analysis reveals functional segregation of release modes and Syt7-dependent facilitation

To definitively separate facilitation from depletion during paired-pulse stimulation, we applied a failure-based analysis that quantifies facilitation under depletion-free conditions. As illustrated in Fig. 6a,b, bouton data were pooled across cells within each genotype and grouped by first-pulse release efficacy, allowing paired-pulse responses to be compared across all trials, trials in which the first action potential failed to evoke release (failure trials), and trials in which it did (success trials). Because failure trials involve no vesicle consumption at the first pulse, they isolate intrinsic facilitation independently of depletion effects (see Methods for implementation details).

**Figure 6.**
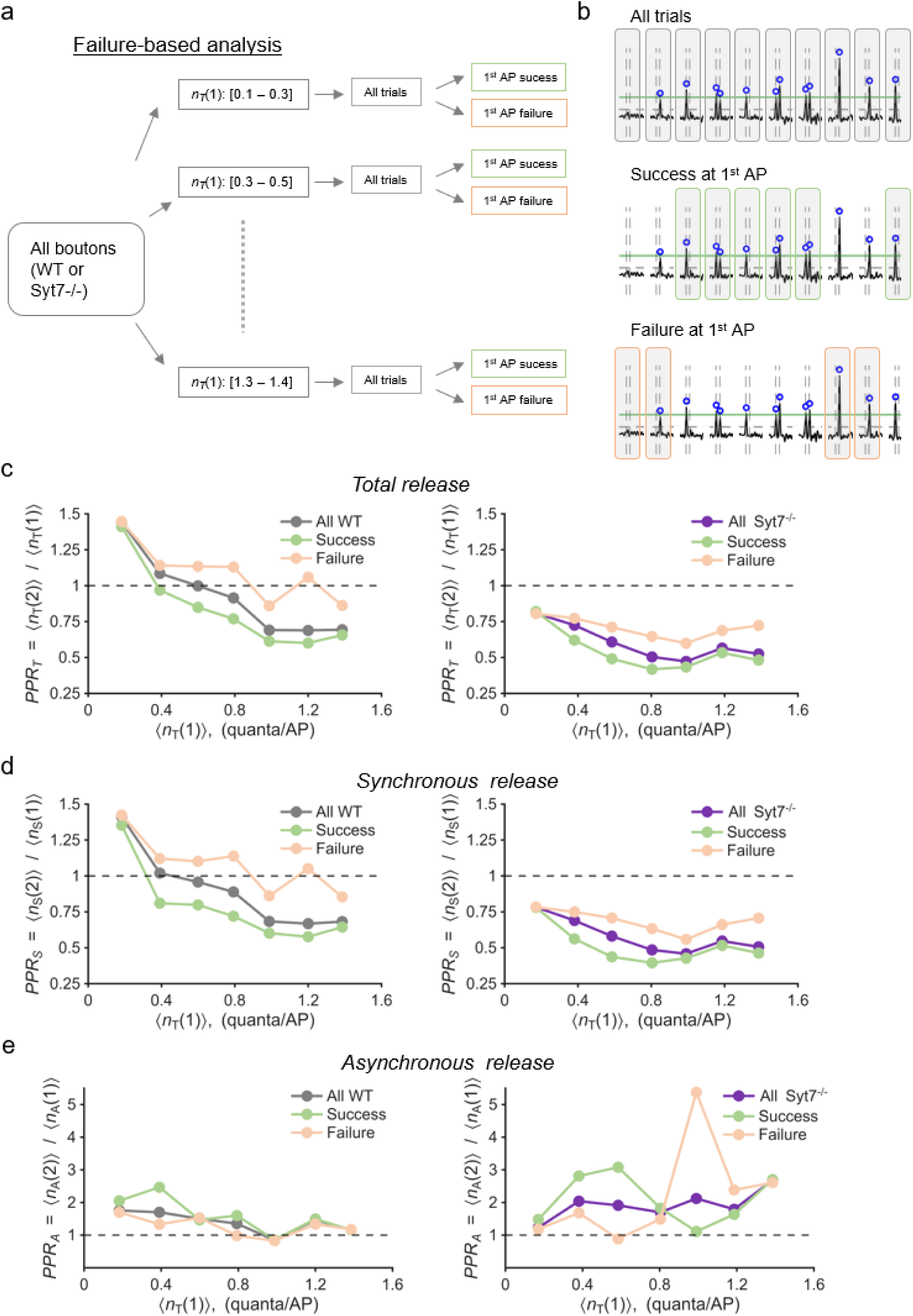
Failure-based analysis reveals functional segregation of release modes and Syt7-dependent facilitation. (a) Schematic of failure-based analysis. Boutons from wild-type (WT) and Syt7^-/-^ neurons were stratified into efficacy bins based on mean first-pulse release *n_T_* (1). Within each bin, paired-pulse ratios were calculated for: all trials 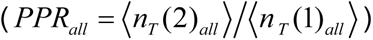, trials where the first action potential evoked release (success subset 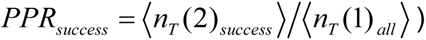, and trials where it failed (failure subset 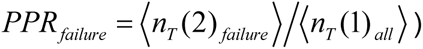. (b) Representative paired-pulse deconvolved SF-iGluSnFR responses from a single WT bouton (*n_T_* (1) bin 0.7 – 0.9). Top: all trials. Middle: success trials (first AP evoked release). Bottom: failure trials (first AP failed to evoke release). Blue circles indicate detected events. (c–e) Paired-pulse ratios as a function of mean 〈*n _T_* (1)*_all_* 〉 for WT (left) and Syt7⁻/⁻ (right) boutons. (c) Total release. (d) Synchronous release. (e) Asynchronous release. Data points represent mean *PPR* within efficacy bins for all trials (grey/purple), success trials (green), and failure trials (orange). Dashed line indicates PPR = 1 (no plasticity). Failure-based *PPRs* represent facilitation under depletion-free conditions, as the first pulse by definition did not consume vesicles. In panel (e), high variance at high efficacies reflects both limited failure trials when release probability is high and the intrinsically low asynchronous release fraction at these boutons. Statistical comparisons in Supplementary Fig. 5.

Analysis of total release revealed clear genotype-dependent differences (Fig. 6c). In wild type neurons, paired-pulse responses during failure trials consistently showed facilitation or absence of depression across the full range of release efficacies, whereas success trials showed a stronger bias toward depression, consistent with vesicle depletion following first-pulse release. At low release efficacies, even success trials exhibited facilitation, but this declined with increasing efficacy and converted to depression at higher efficacies. In contrast, Syt7^-/-^ neurons showed no facilitation in either failure or success trials across the full range of release efficacies. Although paired-pulse ratios during failure trials remained higher than those during success trials, both measures remained below unity, indicating an absence of facilitation even under depletion-free conditions. These results indicate that Syt7 is required for facilitation independently of vesicle depletion.

Applying this same analysis separately to synchronous and asynchronous release components revealed distinct plasticity profiles. Synchronous release (Fig. 6d) closely mirrored total release, with failure trials showing facilitation or no depression in wild type neurons, whereas success trials were shifted toward depression, consistent with vesicle depletion, and these effects were absent in Syt7^-/-^ neurons. In contrast, asynchronous release (Fig. 6e) showed similar facilitation in failure and success trials in both genotypes, with no detectable difference between conditions (Supplementary Fig. 5). This insensitivity of asynchronous facilitation to first-pulse vesicle consumption by the synchronous component, together with its preservation in Syt7^-/-^ neurons, is consistent with facilitation of asynchronous release relying on mechanisms distinct from those that support synchronous facilitation. Together, these findings support functional segregation of synchronous and asynchronous release modes: Syt7 is required for facilitation of synchronous release, but not for facilitation of asynchronous release, indicating that these forms of plasticity being driven by distinct Ca²⁺-dependent pathways.

## Discussion

The calcium sensor Syt7 has been proposed to mediate facilitation, asynchronous release, and vesicle replenishment at central synapses, but its precise contributions remain debated. These roles vary across synapse types, likely reflecting differences in active zone morphology, Ca^2+^ dynamics, and the relative expression of multiple Ca^2+^ sensors including Syt1/2, Syt3, and Doc2α/β (Xu, Mashimo, and Sudhof 2007; Weingarten et al. 2022; Weingarten et al. 2024; Wu et al. 2024; Turecek and Regehr 2019). Using bouton-resolved quantal imaging, we examined how Syt7 shapes release dynamics at individual cortical glutamatergic synapses. We found that Syt7 is essential for facilitation of synchronous release while contributing partially to the magnitude of asynchronous output. Notably, asynchronous facilitation remained intact without Syt7, revealing functionally segregated mechanisms for short-term plasticity of synchronous and asynchronous release.

Single bouton analysis provided critical advantages over population electrophysiology. By imaging 20–180 boutons per neuron with quantal resolution, we directly measured absolute synchronous and asynchronous release efficacies, rather than relying on the asynchronous release fraction, which can be misleading when synchronous and asynchronous components change in parallel. Most importantly, single bouton resolution enabled a failure-based analysis that allowed us to separate facilitation from depletion by examining trials in which the first action potential did not evoke release.

### Syt7 is required for synchronous facilitation independent of depletion

At the cellular level, Syt7^-/-^ neurons showed enhanced depression during paired-pulse and sustained 20 Hz stimulation, confirming that Syt7 supports short-term enhancement (Jackman et al. 2016; Chen et al. 2017; Wu et al. 2024). To determine whether the presence of Syt7 reflects genuine facilitation or simply reduced depletion, we exploited our single trial resolution to perform failure-based analysis. Traditional measurements cannot distinguish increased release probability from reduced depression because successful release inevitably depletes vesicles. By isolating trials in which the first pulse failed, we measured facilitation under depletion-free conditions. In wild type neurons, these failure trials showed robust facilitation of release efficacy or an absence of depression across all efficacy ranges, confirming a true activity-dependent increase in synchronous release probability. In Syt7^-/-^ neurons, facilitation was absent in both failure and success trials at all efficacies, demonstrating that Syt7 directly mediates facilitation rather than merely offsetting depletion. Within the sensitivity of our measurements, alternative mechanisms such as recruitment of other Ca^2+^ sensors or facilitation driven by Ca^2+^ buffer saturation did not generate detectable synchronous facilitation in the absence of Syt7.

### Syt7 contributes to asynchronous release magnitude

The contribution of Syt7 to asynchronous release has been debated, with some studies reporting substantial effects (Turecek and Regehr 2018; Chen et al. 2017; Luo and Sudhof 2017; Bacaj et al. 2013; Weingarten et al. 2024) and others observing minimal changes after Syt7 deletion (Wu et al. 2024). Our single synapse analysis helps to reconcile these discrepancies. At the first action potential, asynchronous release constituted only 3.7% of total release in wild type neurons versus 1.6% in Syt7^-/-^ neurons, a roughly 2-fold reduction that is easily overlooked because at basal conditions asynchronous events represent such a small proportion of total output. Consistent with earlier findings (Mendonca et al. 2022), the asynchronous fraction was highest in low efficacy boutons, but Syt7 deletion reduced asynchronous release by a similar proportion across all bouton types.

This indicates that Syt7 contributes to asynchronous release irrespective of intrinsic release efficacy, while the dominance of high-efficacy synapses in population measurements can mask this effect. Notably, because asynchronous release constitutes a larger fraction of total output at low-efficacy boutons, loss of Syt7 is expected to have a proportionally greater effect on asynchronous output at these synapses.

During 20 Hz trains, measuring Syt7’s contribution required correcting for two confounds: (i) parallel changes in synchronous and asynchronous components that mask absolute reductions when only fractions are examined, and (ii) misclassification of ongoing asynchronous tails as synchronous events. After these corrections, Syt7 deletion reduced absolute asynchronous efficacy by ∼30%, demonstrating consistent but partial contribution to asynchronous release magnitude. In contrast to synchronous release, asynchronous release retained robust facilitation in Syt7^-/-^ neurons during both paired-pulse stimulation and the 20 Hz action potential train, with enhancement ratios that were similar in wild type and Syt7^-/-^ cells. Thus, removal of Syt7 selectively disrupts facilitation of the synchronous pathway while leaving the mechanisms that govern asynchronous facilitation largely intact.

### Functional segregation of synchronous and asynchronous release

Failure-based analysis provided additional evidence for functional segregation of the synchronous and asynchronous pathways. In failure trials paired-pulse ratios for synchronous release were higher both in wild type and Syt7^-/-^ neurons indicating sensitivity to RRP vesicle consumption. In contrast, asynchronous facilitation was identical in the failure and success subsets in both genotypes, showing that it is depletion-insensitive and largely independent of the vesicle population that supports synchronous fusion (that constitutes >95% of release at the first action potential). These observations provide functional evidence that synchronous and asynchronous release occur from at least partially distinct vesicle populations, consistent with partial spatial segregation observed by imaging and electron microscopy (Mendonca et al. 2022; Li et al. 2021; Malagon, Myeong, and Klyachko 2023).

What mechanisms might underlie the functional segregation of synchronous and asynchronous release? One possibility is that RRP vesicles differ in their complement of Ca^2+^ sensors. Vesicles enriched in Syt1/2 would preferentially undergo synchronous fusion, while those with higher Syt7 or other high-affinity sensors could also mediate asynchronous release. Supporting this, reconstitution assays show that Syt1 and Syt7 compete for SNARE binding and increasing Syt7 abundance shifts release kinetics toward asynchronous fusion in a dose-dependent manner (Bose et al. 2024). Alternatively, spatial segregation within active zones could contribute: vesicles near Ca^2+^ channels experience brief, high-amplitude Ca^2+^ nano/micro-domains that activate Syt1/2 for synchronous release, while distant vesicles respond to residual Ca^2+^ via high-affinity sensors for asynchronous fusion (Eggermann et al. 2011; Mendonca et al. 2022).

Our data indicate that in our conditions during paired-pulse protocol Syt7 primarily enhances release probability rather than increasing the RRP size. If Syt7-dependent priming were dominant, failure trials should show facilitation across all efficacy ranges, reflecting expanded RRP independent of basal release probability. Instead, facilitation in wild type neurons was restricted to low/medium-efficacy boutons, while high-efficacy boutons showed minimal facilitation even without depletion. This efficacy-dependent pattern-strong facilitation when initial release probability is low and minimal facilitation when it is already high - is the hallmark signature of a mechanism that enhances release probability rather than expanding vesicle pool size. A pool-expansion mechanism would predict uniform facilitation across all efficacy ranges, as additional vesicles would benefit both low- and high-efficacy synapses equally. The observed pattern is instead consistent with vesicles at high-efficacy boutons already operating near maximal release probability, leaving little room for further enhancement. Additionally, asynchronous facilitation remained intact in Syt7^-/-^ neurons, indicating other high-affinity sensors can also support asynchronous enhancement through residual Ca^2+^ accumulation.

Together with previous studies, our findings support a model where multiple Ca^2+^ sensors have both unique and overlapping roles in regulating synaptic plasticity (Fig. 7). While Syt7 and Syt3 can both promote Ca^2+^-dependent vesicle priming (Wu et al. 2024; Weingarten et al. 2022), and Syt7, Syt3, and Doc2 all contribute to asynchronous release (Weingarten et al. 2024; Bacaj et al. 2013; Yao et al. 2011), Syt7 has a unique function in enhancing the Ca^2+^ sensitivity of synchronous fusion machinery, demonstrated both at different synapses (*e.g.* (Jackman et al. 2016) and this study) and in reconstitution assays (Bose et al. 2024). This model explains why Syt7 deletion specifically abolishes synchronous facilitation while preserving asynchronous facilitation: other sensors cannot compensate for Syt7’s unique role in synchronous release probability enhancement, though they can mediate asynchronous release and its facilitation.

**Figure 7.**
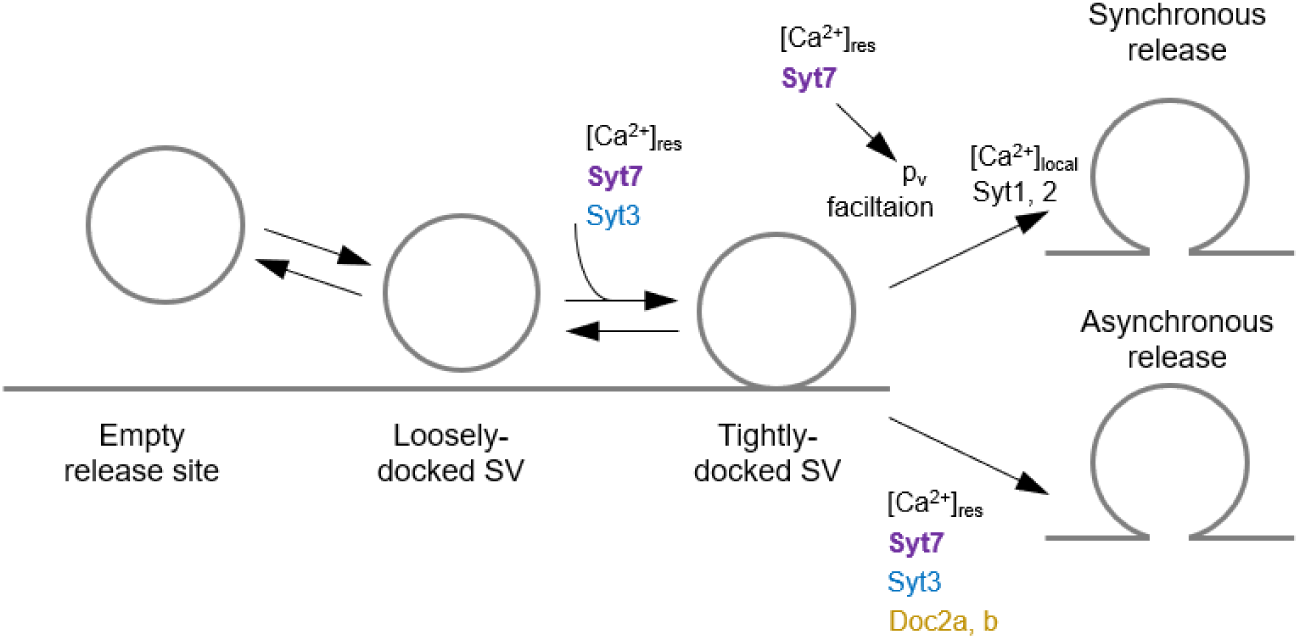
Model for differential regulation of synchronous and asynchronous release by calcium sensors. Vesicles progress from empty release sites through loosely-docked to tightly-docked states in Ca^2+^-dependent manner supported by either Syt7 or Syt3. Tightly-docked vesicles are competent for both synchronous and asynchronous fusion, with the mode determined by Ca^2+^ dynamics and sensor complement. During action potentials, high local [Ca^2+^]_local_ activates low affinity sensors (Syt1/Syt2) for synchronous release, with Syt7 enhancing release probability to enable facilitation. Residual [Ca^2+^]_res_ drives asynchronous release through high-affinity sensors (Syt7, Syt3, Doc2α/β). Syt7 deletion abolishes synchronous facilitation while partially reducing asynchronous release, but asynchronous facilitation persists through remaining high-affinity sensors. This segregation allows independent regulation of synchronous and asynchronous components of synaptic transmission.

## Limitations

Our study focused on Syt7 deletion without manipulating or quantifying other high-affinity Ca²⁺ sensors. Doc2α is present in cultured glutamatergic terminals (Wu et al. 2024), and Syt3 has been reported to be presynaptically expressed in hippocampal pyramidal neurons (Weingarten et al. 2024), though its expression in cortical cultures is unconfirmed. We therefore cannot assess potential interactions between Syt7 and other high-affinity Ca²⁺ sensors in this preparation. A further technical limitation is that all experiments were performed at room temperature (23–25 °C). Previous work at both room and physiological temperatures has reported qualitatively similar forms of short-term plasticity and Syt7-dependent effects, although increasing temperature (32–35 °C) enhances synchronous fusion and reduces the asynchronous release fraction (Huson et al. 2019). Finally, the use of constitutive Syt7 knockout mice leaves open the possibility of developmental compensatory adaptations that cannot be addressed with the present experimental approach.

### Conclusions

Using failure-based analysis enabled by single-bouton resolution, we demonstrate that Syt7 is essential for facilitation of synchronous release at neocortical glutamatergic synapses. In contrast, Syt7 acts alongside other high-affinity Ca^2+^ sensors in supporting asynchronous release and its Ca^2+^-dependent facilitation. This mechanistic segregation indicates that short-term plasticity of synchronous and asynchronous release operates through distinct Ca^2+^-sensing pathways, with important implications for synaptic computation and for the targeting of specific modes of information transfer within neuronal networks.

## Methods

### Neuronal cultures and SF-iGluSnFR expression

Experiments were conducted in accordance with the Animals (Scientific Procedures) Act 1986 and approved by the UK Home Office. Primary cortical neurons were prepared from constitutive Syt7 knockout mice (C57BL/6NTac-Syt7em2(IMPC)H/H, EMMA ID: EM:14671) (Codner et al. 2018), maintained as heterozygous breeding pairs. Neurons from the neocortex of Syt7^-/-^ and Syt7^+/+^ littermates of both sexes (postnatal day 0) were enzymatically dissociated and plated on poly-L-lysine–coated glass coverslips in Neurobasal A medium supplemented with B27 (Thermo Fisher Scientific), as described previously (Mendonca et al. 2022). At 4-6 days in vitro (DIV 4-6), neurons were transfected with the plasmid pAAV.hSynap.SF-iGluSnFR.A184V (Addgene #106174; (Marvin et al. 2018)) using Neuromag reagent (OZ Biosciences). This procedure resulted in sparse expression of SF-iGluSnFR in a small subpopulation of neurons, enabling imaging of individual cells. Experiments were performed between DIV 16 and 21.

### Imaging of glutamate release

SF-iGluSnFR fluorescence imaging was performed on an inverted Axio Observer.A1 microscope equipped with a Prime95B back-illuminated CMOS camera (Teledyne Photometrics) and a 63x oil-immersion objective (1.4 NA). Fluorescence was excited at 470 nm and collected through a 500–550 nm band-pass emission filter. Image acquisition was controlled using μManager software (Edelstein et al. 2014). Experiments were carried out in a custom-built open laminar-flow perfusion chamber (volume 0.35 ml; perfusion rate ∼1 ml min^-1^) maintained at 23–25°C. The extracellular solution contained (in mM): 125 NaCl, 26 NaHCO₃, 12 glucose, 1.25 NaH₂PO₄, 2.5 KCl, 2 CaCl₂, and 1.3 MgCl₂, bubbled with 95% O₂ and 5% CO₂ (pH 7.4). To ensure that recorded SF-iGluSnFR signals originated exclusively from the stimulated neuron, recurrent network activity was blocked by bath application of ionotropic receptor antagonists (in µM): 50 DL-AP5 (Abcam), 10 NBQX (Abcam), and 50 picrotoxin (Tocris Bioscience).

Whole-cell voltage-clamp recordings were obtained from SF-iGluSnFR–expressing pyramidal neurons, following the configuration described in (Mendonca et al. 2022). Recordings were made using a Multiclamp 700B amplifier (Molecular Devices), digitised at 20 kHz and low-pass filtered at 4 kHz. Patch pipettes (4–7 MΩ; Warner Instruments) were filled with an intracellular solution containing (in mM): 125 K^+^ Gluconate, 20 KCl, 10 HEPES, 10 Phosphocreatine-Na_2_, 2 ATP-Mg, 0.4 GTP-NaH_2_0, 0.1 EGTA (pH=7.3, balanced with KOH). Neurons were held at –70 mV, and action potentials were evoked by 5 ms depolarising steps from –70 to –10 mV. This configuration reliably triggered single spikes without burst firing, allowing precise control of presynaptic activity for simultaneous imaging.

After establishing whole-cell recordings, SF-iGluSnFR responses were imaged within the axonal arbour 200–600 µm from the soma at 250 Hz (4 ms per frame). Image acquisition was synchronised with electrophysiological recordings through the data acquisition board, ensuring precise temporal alignment between fluorescence signals and evoked action potentials, as described previously (Mendonca et al. 2022). Between one and six ROIs (1200 x 125 pixels, corresponding to approximately 204 x 22 µm in the specimen plane) were imaged per neuron, with 2–5 min intervals between trials. Each recording comprised nine paired-pulse stimuli at 20 Hz followed by a 50-pulse train at the same frequency, with 10 s between sweeps to allow recovery.

### Image analysis and identification of active boutons

Image analysis was performed offline using ImageJ (NIH) and custom MATLAB scripts, following the procedures described in (Mendonca et al. 2022). Image stacks were first processed with spatial and temporal filters to correct baseline and background fluorescence. A maximal projection of the filtered stack was generated to identify all active boutons that released at least one vesicle during the stimulation protocol. ROIs corresponding to these boutons were automatically selected on the projection image, and fluorescence traces were extracted for further analysis.

Traces were bandpass-filtered (0.5–30 Hz) and deconvolved using the experimentally determined quantal SF-iGluSnFR waveform, characterised by a rapid rise and exponential decay (τ ≈ 68 ms). Deconvolution provided both the timing and amplitude of individual quantal release events.

### Event detection

Event detection thresholds were determined individually for each bouton based on its baseline noise level. Because imaging was performed using the same acquisition protocol and parameters as in (Mendonca et al. 2022), the temporal precision and detection sensitivity of the quantal analysis were identical to those previously established. In that study, the temporal resolution of event detection was limited primarily by the camera acquisition rate (4 ms per frame). Detection thresholds were set individually for each bouton at four times the standard deviation (4σ) of baseline noise, which effectively eliminated false-positive detections. To minimise false negatives, boutons with a signal-to-noise ratio below 5σ were excluded from analysis. During 20 Hz trains, partial temporal overlap between successive events occasionally reduced apparent event amplitude. Although deconvolution resolved the majority of release events, up to approximately 10–15% likely fell below the detection threshold because of overlap or partial indicator saturation. This under-detection may slightly underestimate release efficacy values for the train, but it affects both genotypes equally and therefore does not bias relative comparisons (see Supplementary Figs. 1 and 2). Lowering the detection threshold below four standard deviations of the baseline noise could, in principle, reduce this underestimation; however, as shown previously (Mendonca et al. 2022), this approach substantially increases false-positive detections and artificially inflates the apparent asynchronous component. For this reason, the 4σ threshold was maintained in the present study.

### Quantal analysis

Quantal analysis followed the procedure described in (Mendonca et al. 2022). Amplitude distributions of deconvolved events within individual boutons typically showed discrete peaks corresponding to integer multiples of a quantal response. To obtain stable estimates of quantal amplitude (q) that were insensitive to histogram bin size, amplitude histograms were generated using a bootstrap procedure incorporating bouton-specific noise to produce quasi-continuous distributions. These were then fitted with a mixture of Gaussian functions representing integer multiples of q, with a scaling factor (λ ≈ 0.9) applied to account for partial SF-iGluSnFR saturation during multivesicular release.

Histogram fitting was limited to three Gaussian components, corresponding to the release of one, two, and three vesicles. Inclusion of higher-order Gaussians (four or more vesicles) did not affect the estimated single quantal amplitude but frequently destabilised the fit and introduced artefactual peaks. Rare events with amplitudes consistent with the fusion of four or more vesicles were occasionally observed (for example, WT b3 in Supplementary Fig. 1) but were excluded from fitting to maintain consistency across boutons. This approach yielded robust and unbiased estimates of quantal amplitude and release efficacy parameters across all boutons.

### Calculation of bouton-level functional parameters

From the quantal analysis, three primary parameters were derived for each bouton: total release efficacy (*n_T_*; quanta released per action potential), synchronous release efficacy (*n_S_*), and asynchronous release efficacy (*n_A_*). For the paired-pulse protocol, *n_T_* (1) and *n_T_* (2) denote the mean total number of quanta released per action potential across the ten paired-pulse trials at the first and second stimuli. For the train protocol, *n_T_* (*Tr*) denotes the mean total release efficacy across action potentials 11 to 50 during the train steady-state. Synchronous and asynchronous events were separated using a 10 ms threshold relative to the somatic action potential (see Results for justification). The corresponding synchronous and asynchronous measures, *n_S_* (1), *n_S_* (2), *n_S_* (*Tr*) and *n_A_* (1), *n_A_* (2), *n_A_* (*Tr*), were obtained using the same averaging procedure as for total release. The asynchronous release fraction was expressed as *n_A_*/*n_T_*.

A bouton specific index of short-term synaptic plasticity was calculated either for the second action potential of the paired-pulse protocol or for the steady-state of the stimulus train using the Facilitation-Depression Index (*FDI*):

*FDI_T_* (2,1) = *n_T_* (2)/*mean* [*n_T_*(1), *n_T_* (2)], for the paired-pulse protocol and

*FDI_T_* (2,1) = *n_T_* (*Tr*)/*mean* [*n_T_* (1), *n_T_*(*Tr*)], for the steady-state of the train.

### Adjustment for continued asynchronous release during the steady-state phase of the train

To account for asynchronous release that continues into the next stimulus cycle during the steady-state phase of the train Fig. 3b, we corrected the synchronous and asynchronous components for each bouton. At 20 Hz, each interstimulus interval comprised 12 full frames of 4 ms each (frames 0–11). Frames 0–2 (0–10 ms after the action potential on average; see (Mendonca et al. 2022) for details of the acquisition cycle) were assigned to synchronous release, and frames 3–11 (10–48 ms) to asynchronous release. Because during the steady-state of the train asynchronous release persists across cycles at an approximately constant rate (Fig. 3b), a fraction of asynchronous events from the preceding action potential contaminates the synchronous window of the subsequent cycle. To correct for this, we reduced the synchronous efficacy by the proportion of asynchronous release expected within the 3-frame synchronous window and added this quantity to the asynchronous component:

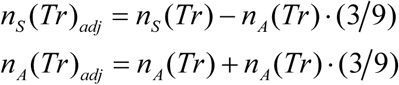

where 3 and 9 are the number of frames in the synchronous and asynchronous windows, respectively. Negative values of *n_S_* (*Tr*)*_adj_* were set to zero, with the corresponding difference added to *n_A_* (*Tr*)*_adj_* to preserve total efficacy. This correction leaves *n_T_* (*Tr*) unchanged and provides a conservative estimate of the true synchronous and asynchronous components during the train steady-state.

### Failure-based analysis of bouton-level facilitation

Bouton data from all cells within each genotype were pooled to ensure continuous sampling across the full range of release efficacies (m = 635 boutons from n = 15 cells for wild-type; m = 933 boutons from n = 19 cells for Syt7^-/-^). Each bouton’s first-pulse release efficacy *n_T_* (1), averaged across ten paired-pulse trials, was used to assign it to fixed efficacy bins (0.1–0.3, 0.3–0.5, …, 1.3–1.5 quanta per action potential). Within each efficacy bin, trial-level data from all boutons were combined into paired vectors of first- and second-pulse responses [*n_T_* (1), *n_T_* (2)].

To calculate paired-pulse ratios while preserving the ability to compare across conditions, we used a common denominator approach. For all trials in each bin, mean 〈*n_T_* (1)*_all_*〉 and mean 〈*n_T_* (2)*_all_*〉 were calculated, with their ratio defining the reference paired-pulse ratio *PPR_all_* = *〈n_T_* (2)*_all_*〉/〈*n _T_* (1)*_all_*〉. The same dataset was then divided into failure (*n_T_* (1) = 0) and success (*n_T_* (1) > 0) subsets. For each subset, mean 〈*n_T_* (2)〉 was computed and divided by the original bin-wide mean 〈*n_T_*(1)*_all_*〉, yielding *PPR_failure_* = 〈*n_T_*(2)*_failure_*〉/〈*n_T_*(1)*_all_* 〉 and *PPR_success_* = 〈*n_T_*(2)*_success_*〉/〈*n_T_*(1)*_all_*〉.

This approach isolates facilitation from depletion effects: failure trials measure facilitation under depletion-free conditions (since no vesicles were consumed by the first pulse), while success trials reflect the net effect of facilitation and depletion. The same analysis was applied separately to synchronous [*n_S_* (1), *n_S_* (2)]. and asynchronous [*n_A_* (1), *n_A_* (2)]. components.

Statistical comparisons across conditions (all, success, failure) within each genotype were performed using the Friedman test for paired data.

### Data Inclusion and Exclusion Criteria

Neurons were included in the analysis only if whole-cell recordings remained stable throughout the imaging session, with access resistance varying by less than 20% and holding current within ±100 pA of the initial value. Cells showing signs of rundown or unstable series resistance were excluded. For cell-level comparisons, all boutons that exhibited at least one detectable release event during paired-pulse stimulations were included, and only neurons with twenty or more active boutons were retained to ensure sufficient sampling. This threshold was based on previously validated criteria (Mendonca et al. 2022), which demonstrated that recordings with approximately twenty boutons per cell provide statistically stable estimates of release parameters across efficacy ranges. For bouton-level analyses, only boutons that allowed meaningful estimation of quantal parameters were included, to avoid division by zero in derived quantities. Specifically, for analysis of the asynchronous release fraction at the first action potential, only boutons with *n_T_* (1) > 0 were included; for analysis of *FDI_T_* (2,1), boutons with *n_T_* (2) + *n_T_* (1) > 0 were included; and for analysis of *FDI_T_* (*Tr*,1), boutons with *n_T_* (*Tr*) + *n_T_* (1) > 0 were included.

### Statistical analysis

Data analysis was performed blind to genotype and condition. All statistical tests were carried out in MATLAB (MathWorks). Detailed results, including test statistics, exact p values, and sample sizes, are provided in Supplementary Table 1 and SourceData.xlsx. No statistical methods were used to predetermine sample sizes; sample sizes were similar to those reported previously for comparable preparations and analyses (Mendonca et al. 2022; Mendonca et al. 2024).

#### Cell-level comparisons

Data distributions for each experimental dataset were first assessed for normality using the Kolmogorov–Smirnov test. None of the measured parameters satisfied normality assumptions, so non-parametric statistics were applied throughout. Between-group comparisons at the level of cells were performed using two-tailed Mann–Whitney U tests, as indicated in figure legends. Summary values are reported as medians with interquartile ranges.

#### Bouton-level comparisons

Because boutons are nested within neurons and bouton counts varied across cells, genotype effects were assessed using linear mixed-effects models (LMMs) to account for within-cell clustering and to avoid pseudoreplication. For each neuron, boutons were sorted by total release efficacy *n_T_* and divided into three equal-sized groups (terciles) corresponding to low-, medium-, and high-efficacy boutons. Within each tercile, mean values were calculated for total release *n_T_* and derived measures (*n_A_/n_T_* or *FDI*).

These per-cell, per-tercile means formed the basis of the hierarchical analyses.

To characterise how each parameter depended on release efficacy within a genotype, we first fitted Model 1 (efficacy dependence):

Outcome_tercile ∼ *n_T_*_tercile + (1 | CellID)

To test whether the dependence on release efficacy differed between genotypes, we then fitted Model 2 (genotype by efficacy):

Outcome_tercile ∼ *n_T_*_tercile × Genotype + (1 | CellID)

Models were fitted by restricted maximum likelihood (REML) in MATLAB (fitlme), and inference was based on the Wald t statistics returned by fitlme. Model coefficients, confidence intervals, and exact p values are provided in SourceData.xlsx.

#### Failure-based analysis

Statistical evaluation of success–failure differences within the failure-based analysis (Fig. 6; Supplementary Fig. 5) was performed using the Friedman test, the non-parametric equivalent of repeated-measures ANOVA. When significant main effects were detected, post-hoc pairwise comparisons were conducted using the Tukey–Kramer procedure on ranked data (MATLAB multcompare function). Exact p values and post-hoc comparisons are reported in SourceData.xlsx.

Because the failure-based analysis was designed to examine facilitation as a function of bouton release efficacy, data were pooled at the bouton level within each genotype to ensure continuous sampling across the full efficacy range. This population-level approach does not model cell-to-cell variability and therefore cannot be interpreted as a hierarchical test across neurons. However, it provides a direct measure of release dynamics at the level of individual synapses, which is the relevant unit for assessing facilitation and depletion. The consistency of trends across independent cells and the close correspondence with cell-level analyses support the robustness of this approach (Figs. 2 and 5).

## Data availability

Source data underlying the figures will be made publicly available upon publication.

## Code availability

Custom MATLAB code for quantal analysis was used as described in (Mendonca et al. 2022) and is available for download from that publication.

## Acknowledgments and funding

We thank Drs Dimitri Kullmann for critical feedback and James Rothman for access to resources and materials. The work was funded by The Wellcome Trust Strategic Award 104033/z/14/z (K.E.V.), The Wellcome Trust PhD Studentship 203795/Z/16/Z (H.L. and K.E.V.); National Institute of Health (NIH) grant NS133091 (S.S.K. and K.E.V.); UKRI MRC Project Grant MR/T002786/1 (Y.T. and K.E.V.)

## Author contributions

Conceptualisation: Y.T., S.S.K. and K.E.V. Performing experiments: D.K., H.L. and E.MG. Development of methodology and data analysis framework: D.K., H.L., P.R.F.M., E.T., Y.T. and K.E.V. Writing original draft: K.E.V. All authors discussed the results and commented on the manuscript. Funding acquisition: H.L., Y.T., S.S.K., and K.E.V.

## Supplementary materials

**Supplementary Table 1.**
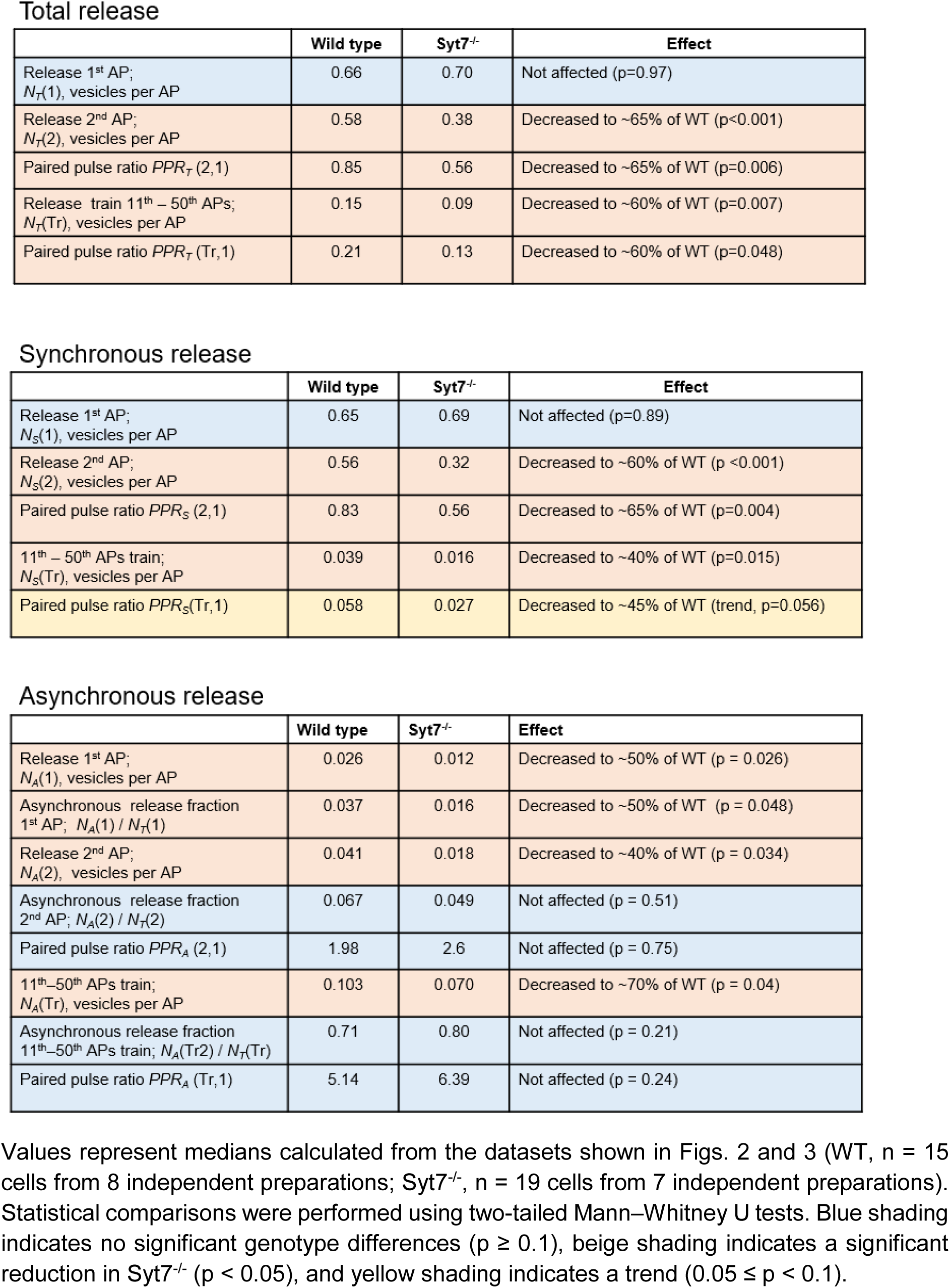
Median cell-level release parameters in wild type and Syt7^-/-^ neurons during paired-pulse and train stimulation.

**Supplementary Figure 1.**
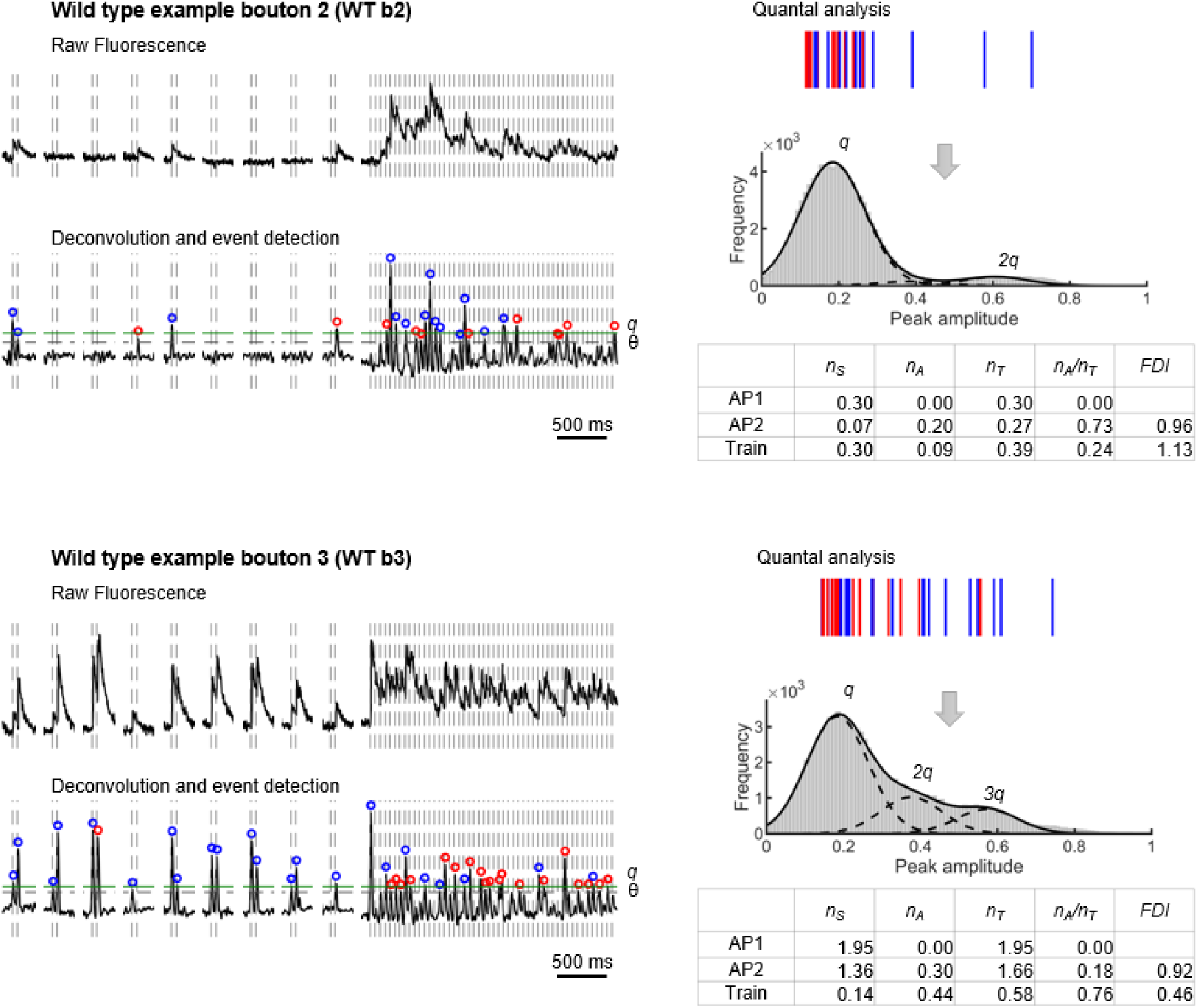
Quantal analysis in representative wild type boutons (related to Fig. 1). Two additional representative wild type boutons (WT b2 and WT b3) analysed using the same quantal imaging and analysis pipeline as in Fig. 1. Left: raw fluorescence traces (top) and corresponding deconvolved signals (bottom). Vertical dotted lines indicate action potential (AP) timings; gaps correspond to 10 s intersweep intervals. The detection threshold (θ) is shown as a horizontal dashed line. Blue and red circles mark synchronous and asynchronous release events (<10 ms and >10 ms after the preceding AP, respectively). Right: amplitude distributions of deconvolved events were converted into quasi-continuous histograms using a bootstrap procedure incorporating bouton-specific noise and fitted with Gaussian mixtures to estimate the quantal amplitude (q). Peaks at q, 2q, and 3q correspond to the release of one, two, or three vesicles. Gaussian fitting was restricted to three components to ensure stable estimation of q (see Methods). Rare higher-amplitude events consistent with the release of four or more vesicles were occasionally observed (for example, WT b3) but were not included in the fitting procedure. Tables summarise bouton-specific quantal parameters for the first (AP1) and second (AP2) pulses of the paired-pulse protocol and for the steady-state phase of the train (Train; APs 11– 50). Partial overlap of successive events during 20 Hz trains occasionally reduced apparent event amplitudes, leading to minor under-detection (∼10–15%) of small asynchronous events (see Methods).

**Supplementary Figure 2.**
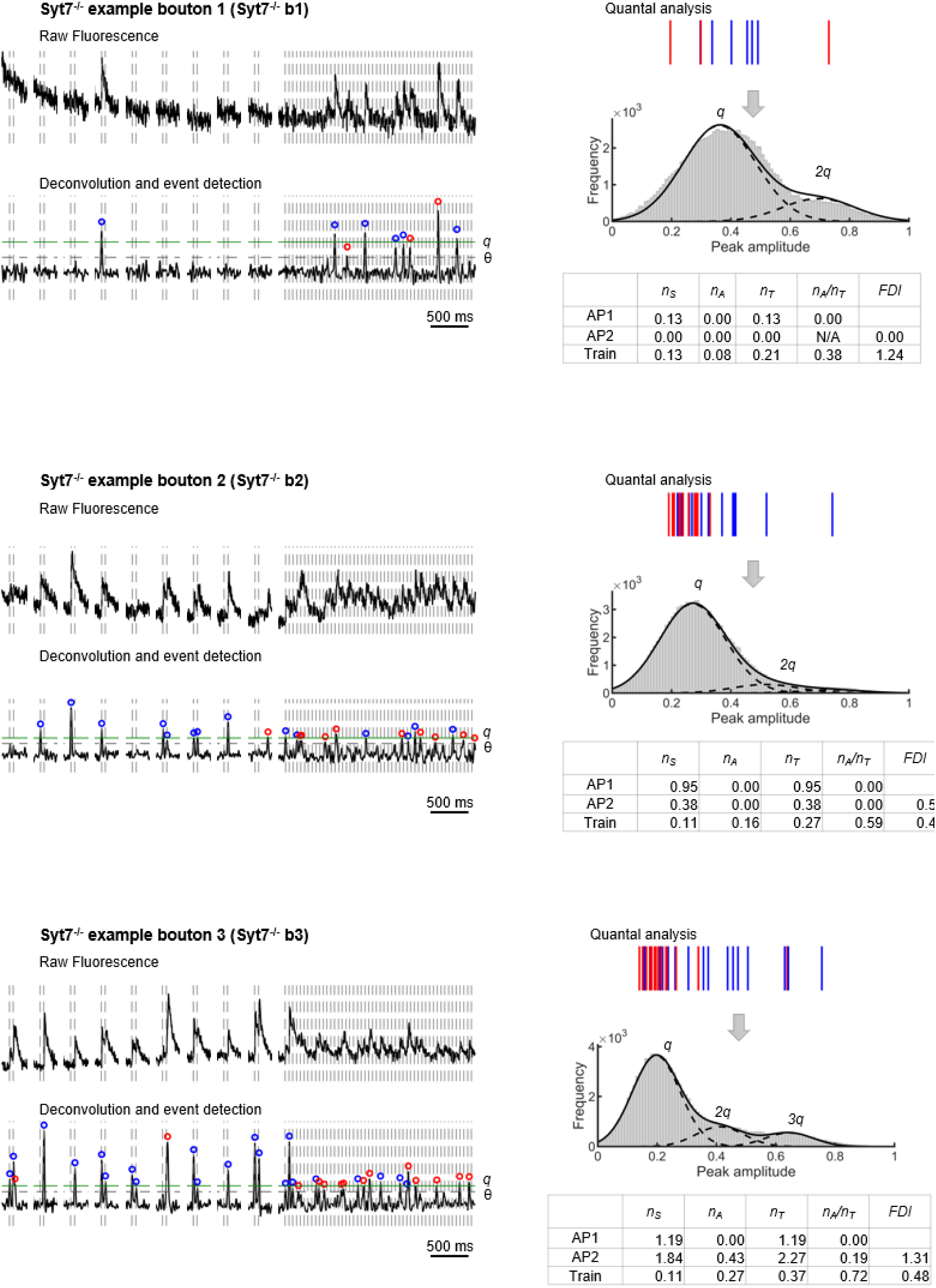
Quantal analysis in representative Syt7^-/-^ boutons (related to Fig. 1) Three representative boutons from Syt7⁻/⁻ neurons (Syt7⁻/⁻ b1–b3) analysed using the same quantal imaging and analysis pipeline as in Fig. 1. Left: raw fluorescence traces (top) and corresponding deconvolved signals (bottom). Vertical dotted lines indicate action potential (AP) timings; gaps correspond to 10 s intersweep intervals. The detection threshold (θ) is shown as a horizontal dashed line. Blue and red circles mark synchronous and asynchronous release events (<10 ms and >10 ms after the preceding AP, respectively). Right: amplitude distributions of deconvolved events were converted into quasi-continuous histograms using a bootstrap procedure incorporating bouton-specific noise and fitted with Gaussian mixtures to estimate the quantal amplitude (q). Peaks at q, 2q, and 3q correspond to the release of one, two, or three vesicles. Gaussian fitting was restricted to three components to ensure stable estimation of q (see Methods); in these examples, no events exceeded 3q. Tables summarise bouton-specific quantal parameters for the first (AP1) and second (AP2) pulses of the paired-pulse protocol and for the steady-state phase of the train (Train; APs 11– 50). Partial overlap of successive events during 20 Hz trains occasionally reduced apparent event amplitudes, leading to minor under-detection (∼10–15%) of small asynchronous events (see Methods).

**Supplementary Figure 3.**
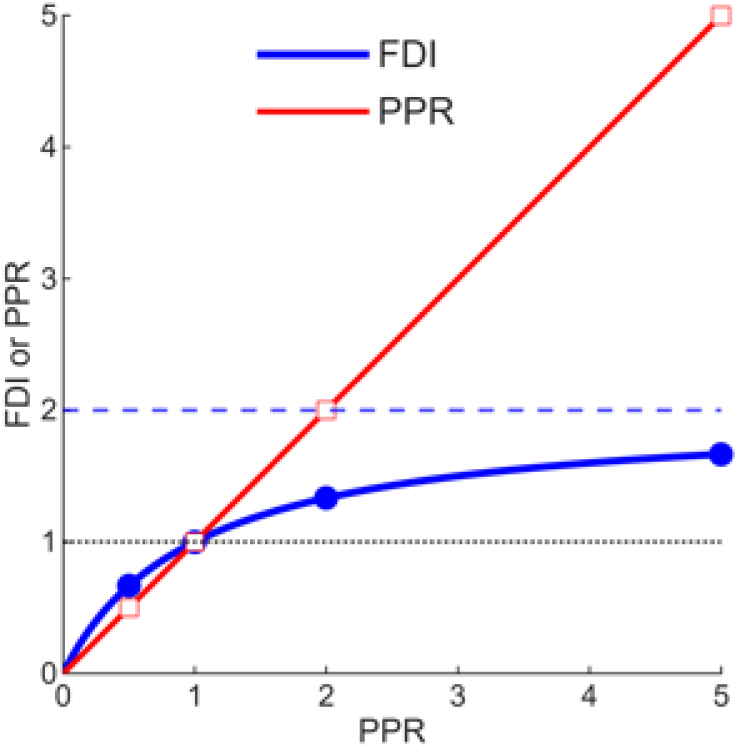
Comparison of conventional paired-pulse ratio (PPR) and Facilitation-Depression Index (FDI) definitions. Relationship between *PPR_T_* (2,1) = *n_T_* (2)*/n_T_* (1) (red line) and *FDI_T_* (2,1) = *n_T_* (2)/*mean* [*n_T_* (1), *n_T_* (2)] (blue line). The two measures are related by 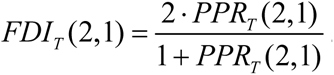. Both measures coincide when equal to 1 (dotted black line) and yield comparable values for depression (*PPR* < 1). Under facilitation (*PPR* > 1), *FDI_T_* (2,1) approaches an upper limit of 2 (dashed blue line) because it is normalised to the mean of the first and second responses. This formulation avoids division by zero in boutons that fail to release across all paired-pulse trials at the first action potential, enabling consistent quantification of short-term plasticity across boutons with a wide range of release efficacies.

**Supplementary Figure 4.**
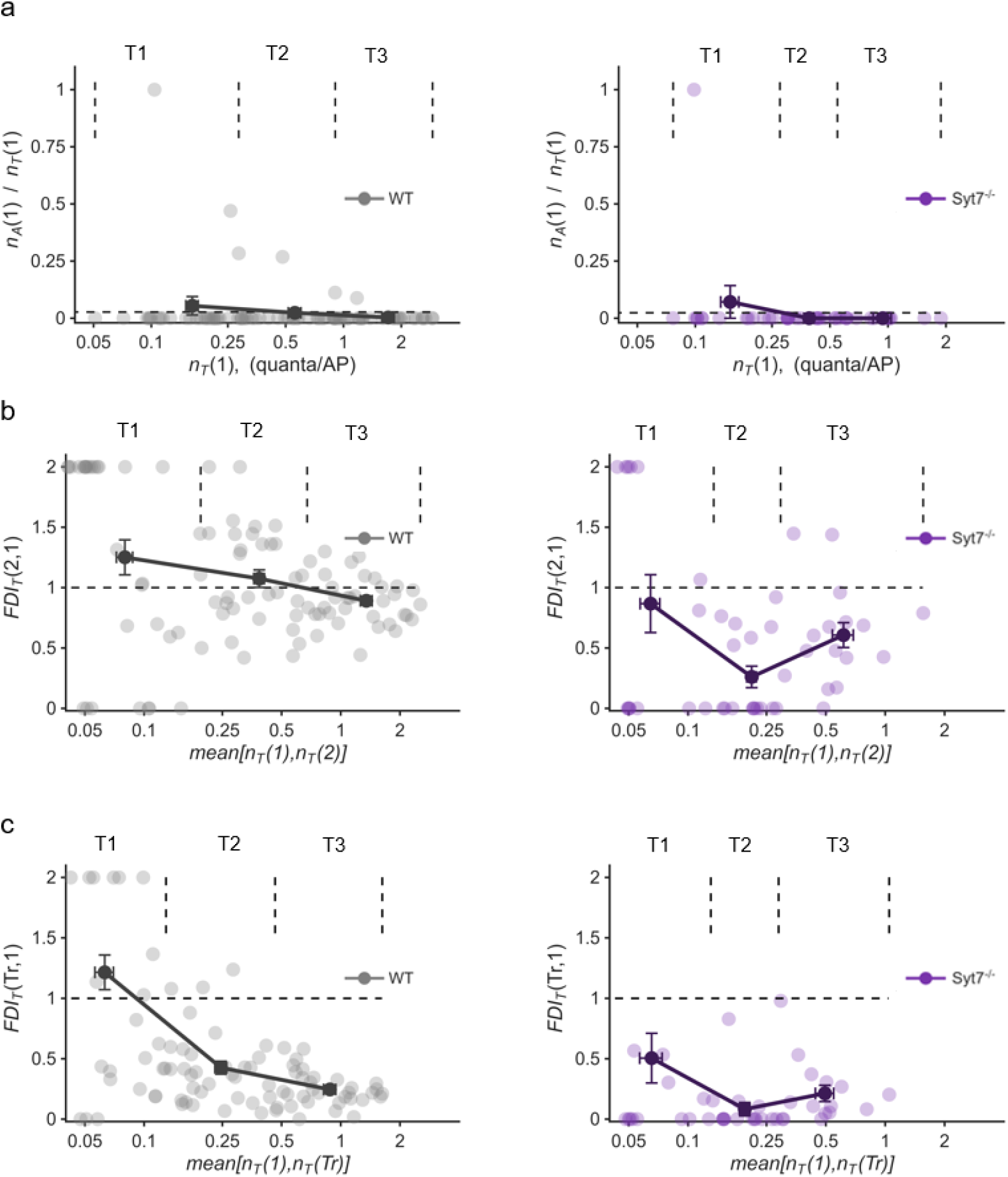
Bouton-to-bouton heterogeneity of release properties within single neurons. Representative examples from one wild type neuron (n = 101 boutons) and one Syt7^-/-^ neuron (n = 48 boutons) illustrating variability of release properties across boutons supplied by the same axon. Boutons were ranked by total release efficacy and divided into three equal-sized groups (terciles), as in Figs. 4 and 5. Shown are relationships between (a) the asynchronous release fraction at the first action potential *n_A_* (1)/*n_T_* (1) and total release efficacy *n_T_* (1); (b) short-term plasticity during paired-pulse stimulation, quantified using the Facilitation-Depression Index *FDI_T_* (2,1), plotted against the mean total release across the first and second action potentials; and (c) short-term plasticity during the steady-state phase of the stimulus train, quantified as *FDI_T_* (*Tr*,1), plotted against the mean total release across the first and steady-state responses. Vertical dashed lines indicate the division of boutons into low-(T1), medium-(T2), and high-efficacy (T3) terciles. Dark-coloured points connected by lines show tercile means ± SEM within each neuron. Dashed horizontal lines in (a) indicate the cell-averaged asynchronous release fraction, and dashed horizontal lines in (b, c) indicate *FDI* = 1, corresponding to no facilitation or depression. The number of boutons contributing to each panel differs due to stimulus-specific inclusion criteria (see Methods, Data Inclusion and Exclusion Criteria).

**Supplementary Figure 5.**
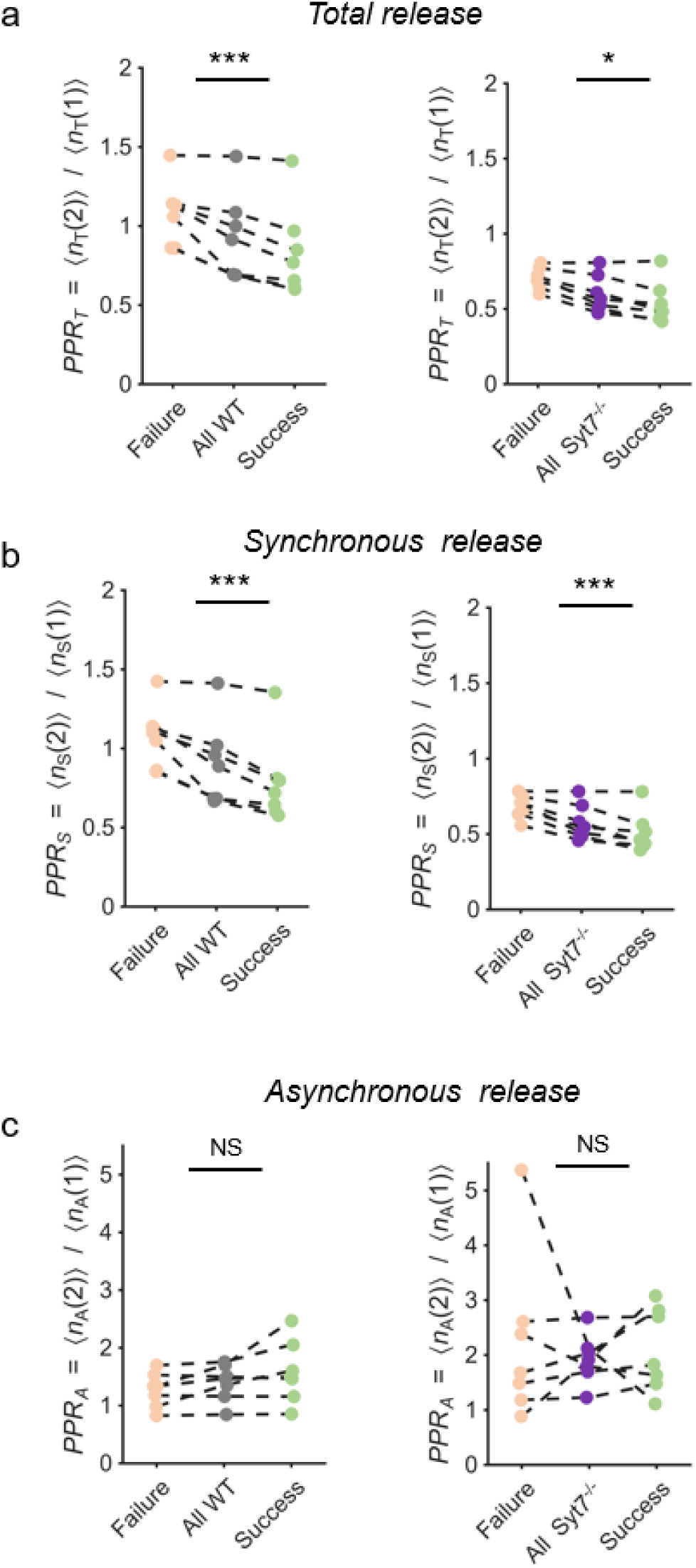
Paired-pulse ratios for total and synchronous, but not asynchronous, release depend on first-pulse outcome. (a–c) Paired comparisons from the failure-based analysis in Fig. 6. For each genotype, paired-pulse ratios were computed separately for three trial subsets: all trials, trials in which the first action potential evoked release (success), and trials in which it did not (failure). Within each first-pulse efficacy bin, these three values form a matched set, and statistical significance was assessed using the Friedman test. Dashed lines connect matched data points from the same efficacy bin across the three conditions. Left column, wild type; right column, Syt7⁻/⁻. (a) Total release (*PPR_T_*). (b) Synchronous release (*PPR_S_*). (c) Asynchronous release (*PPR_A_*). For total and synchronous release, the distribution of paired-pulse ratios differed significantly across failure, all-trial, and success conditions in both genotypes (Friedman test; *p < 0.05, ***p < 0.001). In contrast, asynchronous release showed no significant difference across conditions in either genotype (NS). These results indicate that the paired-pulse ratio of the asynchronous component, unlike that of total and synchronous release, is unaffected by whether the first stimulus evoked release.

